# Juvenile exposure to acute traumatic stress leads to long-lasting alterations in grey matter myelination in adult female but not male rats

**DOI:** 10.1101/2020.12.14.422686

**Authors:** Jocelyn M. Breton, Matthew Barraza, Kelsey Y. Hu, Samantha Joy Frias, Kimberly L.P. Long, Daniela Kaufer

## Abstract

Stress early in life can have a major impact on brain development, and there is increasing evidence that childhood stress confers vulnerability for later developing psychiatric disorders. In particular, during peri-adolescence, brain regions crucial for emotional regulation, such as the prefrontal cortex (PFC), amygdala (AMY) and hippocampus (HPC), are still developing and are highly sensitive to stress. Changes in myelin levels have been implicated in mental illnesses and stress effects on myelin and oligodendrocytes (OLs) are beginning to be explored as a novel and underappreciated mechanism underlying psychopathologies. Yet there is little research on the effects of acute stress on myelin during peri-adolescence, and even less work exploring sex-differences. Here, we used a rodent model to test the hypothesis that exposure to acute traumatic stress as a juvenile would induce changes in OLs and myelin content across limbic brain regions. Male and female juvenile rats underwent three hours of restraint stress with exposure to a predator odor on postnatal day (p) 28. Acute stress induced a physiological response, increasing corticosterone release and reducing weight gain in stress-exposed animals. Brain sections containing the PFC, AMY and HPC were taken either in adolescence (p40), or in adulthood (p95) and stained for markers of OLs and myelin. We found that acute stress induced sex-specific changes in grey matter (GM) myelination and OLs in both the short- and long-term. Exposure to a single stressor as a juvenile increased GM myelin content in the AMY and HPC in p40 males, compared to the respective control group. At p40, corticosterone release during stress exposure was also positively correlated with GM myelin content in the AMY of male rats. Single exposure to juvenile stress also led to long-term effects exclusively in female rats. Compared to controls, stress-exposed females showed reduced GM myelin content in all three brain regions. Acute stress exposure decreased PFC and HPC OL density in p40 females, perhaps contributing towards this observed long-term decrease in myelin content. Overall, our findings suggest that the juvenile brain is vulnerable to exposure to a brief severe stressor. Exposure to a single short traumatic event during peri-adolescence produces long-lasting changes in GM myelin content in the adult brain of female, but not male, rats. These findings highlight myelin plasticity as a potential contributor to sex-specific sensitivity to perturbation during a critical window of development.

## 1. Introduction

Stress early in life can have a major impact on brain development and behavior. In particular, stressful experiences from infancy through adolescence are associated with an increased risk of later developing psychiatric disorders (Carr, Martins, Stingel, Lemgruber, & Juruena, 2013; Hughes et al., 2017; Ventriglio, Gentile, Baldessarini, & Bellomo, 2015). For example, childhood trauma increases the risk for developing depression and anxiety (Heim & Nemeroff, 2001). Yet, individuals who experience similar stressful environments can have very different responses to stress; only a subpopulation demonstrates vulnerability, while others demonstrate resilience (Compas & Phares, 1991; Kessler, Sonnega, Bromet, Hughes, & Nelson, 1995; McEwen & Stellar, 1993). In part, these individual differences may be explained by biological sensitivities to context and the environment (Ellis & Boyce, 2008). Furthermore, other factors, such as sex, may play a role. For example, females are more susceptible to developing post-traumatic stress disorder (PTSD) and anxiety (Breslau, 2009; Kessler et al., 1995; McLean & Anderson, 2009). Thus, a major goal is not only to understand the neurobiological effects of early life stress, but also to understand the biological factors that contribute to individual variability.

Experiencing early life stress, encompassing infancy through adolescence, leads to physiological changes in the body and in the central nervous system (Bolton, Short, Simeone, Daglian, & Baram, 2019; Pagliaccio et al., 2014; Van Bodegom, Homberg, & Henckens, 2017). In particular, stressors experienced during peri-adolescence may have a significant impact on brain maturation and development. Adolescence, defined by the onset of puberty (~age 10 in humans, ~postnatal day 36 in rodents), is a major period of experience-dependent plasticity and thus, the brain is particularly sensitive to environmental stimuli such as stressors (Piekarski et al., 2017). This in turn could contribute towards the onset of psychiatric disorders such as anxiety and depression, which often appear around this time (Eiland & Romeo, 2013; Gee & Casey, 2015; Kessler et al., 2007, 2005). Importantly, brain regions that play a role in the stress response, such as the amygdala (AMY), hippocampus (HPC) and prefrontal cortex (PFC), are still developing during periadolescence and are highly sensitive to stress (McEwen et al., 2015; Popoli, Yan, McEwen, & Sanacora, 2012; Roozendaal, McEwen, & Chattarji, 2009; Spear, 2000).

In humans, stress early in life leads to alterations in multiple brain regions, including in the HPC, AMY and PFC. For example, early life stress reduced hippocampal and amygdala volume and led to alterations in the frontal cortex and anterior cingulate cortex (Andersen et al., 2008; R. A. Cohen et al., 2006; Hanson et al., 2013; Luby et al., 2013; Teicher, Samson, Polcari, & McGreenery, 2006). In one specific longitudinal study of children who experienced maltreatment, high cortisol levels and PTSD symptoms were correlated with subsequent reductions in hippocampal volume (Carrion, Weems, & Reiss, 2007). In addition to structural changes in grey matter, early life stress also changes functional connectivity. In particular, there is elevated amygdala reactivity to emotional stimuli and weaker amygdala – PFC connectivity after early stress (Gee et al., 2013; McCrory et al., 2013; Nooner et al., 2013; Tottenham & Galván, 2016). In rodents, structural and connectivity changes in the hippocampus, amygdala and PFC are also observed following early life stress (M. M. Cohen et al., 2013; Gutman & Nemeroff, 2002; Honeycutt et al., 2020; Johnson et al., 2018; Meaney, Aitken, Van Berkel, Bhatnagar, & Sapolsky, 1988). While the majority of rodent studies have utilized early postnatal stressors, resembling infancy in humans, later early-life periods such as the juvenile and/or peri-adolescent period, resembling childhood and pre-teen years in humans, have only recently been explored (Eiland, Ramroop, Hill, Manley, & McEwen, 2012; Eiland & Romeo, 2013; Isgor, Kabbaj, Akil, & Watson, 2004; Oztan, Aydin, & Isgor, 2011; Spear, 2000). In 2012, Eiland and colleagues found that chronic restraint stress during the periadolescent period (p20-41) in male and female rats reduced pyramidal neuron complexity in the PFC and hippocampus but increased neuronal complexity in amygdala. Furthermore, these changes were associated with elevated depressive-like behaviors (Eiland et al., 2012). Thus, in both humans and rodents, there is a growing body of literature suggesting that stress exposure throughout early life leads to changes in developing limbic brain regions, including the hippocampus, amygdala and PFC.

Much of the peri-adolescent literature has focused on the effects of stress on neurons and neuronal plasticity. However, stress effects on glia are beginning to be explored. Specifically, oligodendrocytes (OLs) and the myelin they produce are sensitive to stress, not only in white matter, but also in grey matter regions (Chetty et al., 2014; Makinodan, Rosen, Ito, & Corfas, 2012; Monje, 2018; Saul, Helmreich, Rehman, & Fudge, 2015). In addition, OLs and myelin have been implicated in a number of mental health disorders, including schizophrenia, depression and PTSD, suggesting they play a functional role in mood (Birey, Kokkosis, & Aguirre, 2017; Chao, Tosun, Woodward, Kaufer, & Neylan, 2015; Falkai et al., 2016; Fields, 2008; Lee & Fields, 2009; Ma et al., 2007; Nave & Ehrenreich, 2014; Regenold et al., 2007; Sokolov, 2007; Tham, Woon, Sum, Lee, & Sim, 2011). Chronic stress early in life can also alter myelination in both humans and rodents. In humans, both institutionalization and early child abuse are associated with alterations in white matter in the PFC; furthermore, these changes correlate with cognitive deficits (Hanson et al., 2013; Lutz et al., 2017). In mice, chronic social isolation during peri-adolescence leads to increased depressive-like behaviors, reduced myelin basic protein (MBP) and hypomyelination of the PFC (Leussis & Andersen, 2008; Makinodan et al., 2012). This effect is only observed when the stressor occurs during the juvenile period, following weaning but prior to puberty, suggesting there may be a critical window for stress effects on PFC myelination (Makinodan et al., 2012). Together, these data indicate that altered myelination may be a novel and underappreciated mechanism by which psychopathologies emerge.

While the majority of prior work has focused on chronic stress effects on myelin and OLs, less is known about the effects of acute traumatic events. Experiencing trauma in childhood leads to increased risk for developing psychiatric disorders, including anxiety, depression and PTSD; these same disorders are also associated with alterations in OLs and myelin (Fields, 2008; Heim & Nemeroff, 2001; Yehuda, Halligan, & Grossman, 2001). A critical unanswered question then is whether changes in OLs and myelin are observed following acute traumatic stress during the juvenile period. In particular, the PFC, AMY and HPC are key limbic brain regions of interest, as they are highly plastic during peri-adolescence and chronic stress robustly alters myelin across these regions. Thus, in this study, we sought to explore whether experiencing acute stress during peri-adolescence will induce changes in OLs and grey matter (GM) myelin content across limbic regions involved in stress and emotional regulation. Furthermore, we aimed to assess both short and long-term consequences. As peri-adolescence is a period of heightened experience-dependent plasticity, we predicted that acute stress would result in altered myelination in developing limbic brain regions. Specifically, in line with the chronic stress literature, we hypothesized that there would be decreased myelination following acute stress. Lastly, little is known about how myelin and OLs relate to individual differences following trauma. Therefore, we sought to address if physiological responses to traumatic stress were associated with myelin and OLs in these regions. In addition, we examined whether there are sex differences in these measures following exposure to acute severe stress. Sex is an important biological factor that contributes to individual variation in response to stress. To test these questions, we first exposed male and female juvenile rats to an acute, severe stressor. We then analyzed OL and myelin markers in the PFC, AMY and HPC in order to examine the effects of stress on glial plasticity. Tissue was taken either from adolescent or adult animals in order to test for short- or long-term changes respectively. Lastly, we assessed whether myelin and OL markers correlated with corticosterone responses throughout stress exposure, with the hypothesis that animals with the greatest physiological changes would also display the greatest changes in myelin, relative to controls.

## 2. Methods

### 2.1. Animals

Sixty-four male and female Sprague Dawley rats were used for these experiments. All rats were bred in-house in order to minimize stressful experiences such as shipping prior to testing. Rats were weaned at p21, pair-housed, given *ad libitum* access to food and water and kept on a 12/12 hr. light/dark cycle. All procedures were approved by UC Berkeley’s Animal Care and Use Committee.

### 2.2. Stress

Each cage of rats was randomly assigned to either a stress or control condition. For animals in the stress condition (n=32), at postnatal day 28 (p28), juvenile male and female Sprague Dawley rats were exposed to three hours of severe stress (immobilization with exposure to a predator odor; n=16 males, n=16 females; Figure 1A). Specifically, rats were restrained in plastic Decapicone bags (Braintree Scientific, Inc, Braintree, MA) and placed in a clean cage with a predator odor inside a fume hood. The cage contained a cotton ball infused with 1mL of synthetic fox urine (Red Fox Urine, Trap Shack Company, Arcadia, WI) taped approximately 1 inch from the animal’s nose. Cagemates were placed side by side in the cage for the extent of the stressor. Blood samples from the tail vein were collected at three time points (see details below). All stress testing was conducted between the hours of 8am and noon. At the end of the three-hour stressor, cagemates were returned to a clean cage and allowed to self-groom. All animals in the stress condition were kept in a separate housing room for three days prior to being returned to their normal housing room, in order to minimize stress transmission to other rats.

**Figure 1.**
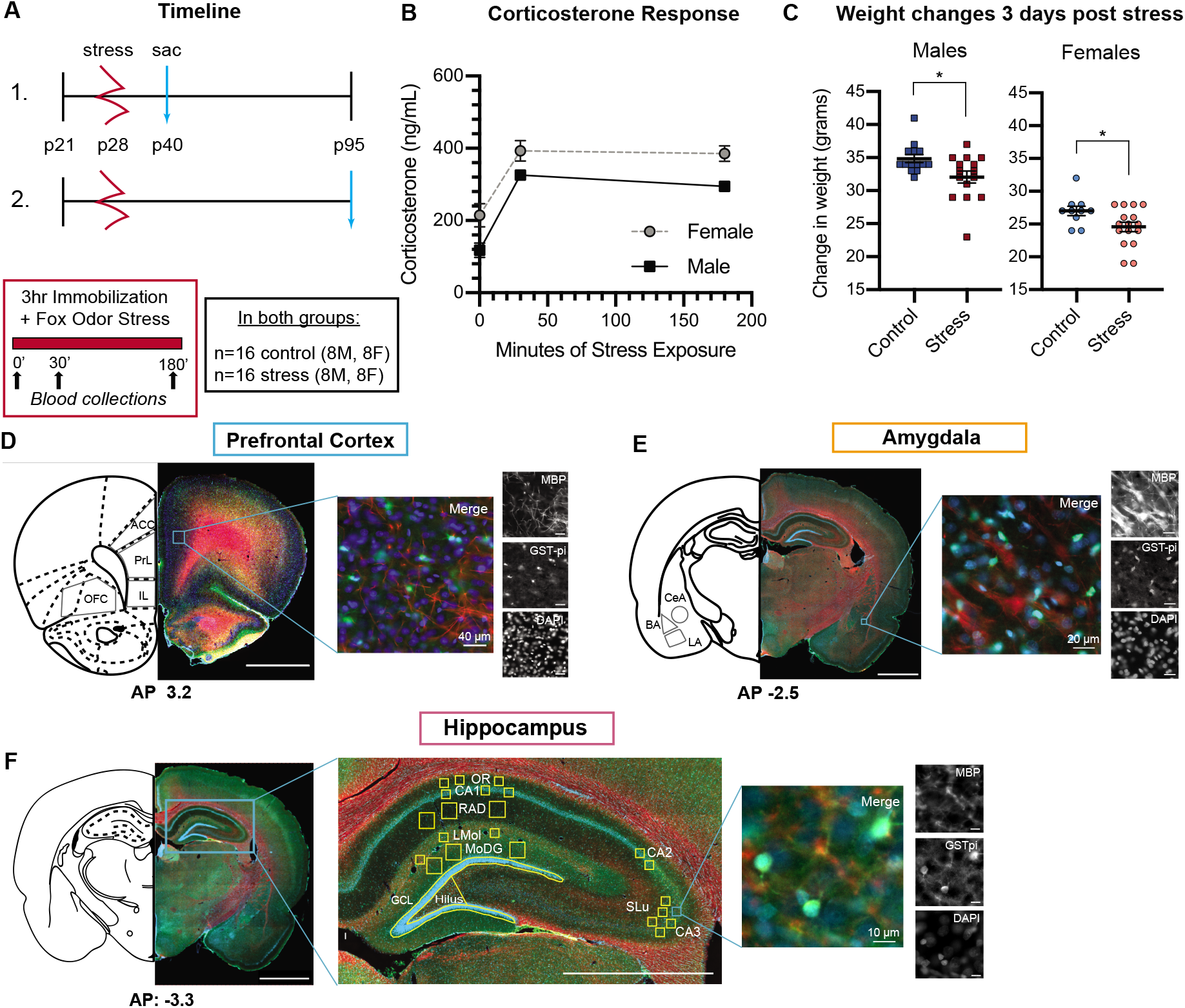
Methods and Physiological responses to acute traumatic stress. **A)** Experimental Timeline. Animals were divided into two cohorts: one tested in the short term (1) and one tested in the long term (2). For both cohorts, animals underwent three hours of immobilization stress with exposure to fox urine (stress) at postnatal day 28 (p28). (n=8 males, n=8 females in each cohort). Blood was collected at three timepoints: just prior to stress (0’), 30 minutes into the stress (30’), and at the end of stress at 180 minutes (180’). An additional control group remained in their home cage (n=8 males, n=8 females in each cohort). One cohort of animals was sacrificed (sac) at p40, while another cohort was sacrificed at p95 when they were adults. **B)** Corticosterone responses to acute traumatic stress. Corticosterone levels (ng/mL) robustly increased during exposure to the stressor for both male and females, with higher corticosterone at 30 and 180 minutes over baseline values. **C)** Weight changes three days post stress for animals in both cohorts. Statistically significant differences are marked with asterisks (*p<0.05). **D-F:** Example prefrontal cortex (PFC), amygdala (AMY) and hippocampal (HPC) coronal sections stained for MBP (red), GST-pi (green) and DAPI (blue). All coordinates are anterior-posterior (AP) from bregma. **D)** Example PFC section. Subregions analyzed are labeled in grey. Scale bar 2mm. On right: Representative staining in the prelimbic cortex. **E)** Example AMY section. Subregions analyzed are labeled in grey. Scale bar 2mm. On right: Representative staining in the basal amygdala. **F)** Example HPC section. Scale bar 2mm. Center: A zoom in on the hippocampus. Subregions analyzed are labeled in yellow. Scale bar 1mm. On right: Representative staining in the CA3 subregion.

### 2.3. Weight collections

Animals in both conditions were weighed at p28. Animals in the stress condition were weighed immediately prior to placement in the stress manipulation, while animals in the control condition (n=32, 16 males, 16 females) were weighed in the housing room and otherwise remained in their home cage undisturbed. On the day stress animals were returned to their original housing room (three days post-stress exposure), rats in all conditions were weighed again. Weights from six female control animals were excluded in analyses due to inaccurate measurements.

### 2.4. Serum sampling for corticosterone analysis

Once restrained, tail blood was collected from each rat at 0 minutes, 30 minutes into, and at the end (3 hours) of the acute traumatic stressor. Specifically, a sterile scalpel was used to remove a small segment at the end of the tail, and approximately 0.1-0.2mL of blood was collected at each time point. For the baseline time point at the start of the stressor, we collected blood within the first two minutes to avoid detecting elevations in corticosterone due to the restraint itself. All collected samples were kept on ice throughout the stressor. Blood clots were then removed, and samples were centrifuged at 9,391 g for 20 minutes at 4°C. Serum was then extracted and stored in clean tubes at −80°C. Samples were assayed using a Corticosterone EIA kit (Arbor Assays, Ann Arbor, MI), with 2 replicates per sample. Samples were compared against a standard curve to obtain the concentration of corticosterone within each sample. Using corticosterone values from the three time points throughout stress exposure (0, 30 and 180 minutes), we calculated an area under the curve (AUC) for each animal. This provides us with an overall measure of the corticosterone response.

### 2.5. Perfusions and Brain Extractions

Rats were euthanized either at p40 (n=32; 16 control, 16 stress) or p95 (n=32, 16 control, 16 stress) to test for short- or long-term effects, respectively. Animals were weighed, then deeply anesthetized with sodium pentobarbital 200 mg/kg (Euthasol ^®^, Vibrac AH Inc.) and transcardially perfused with ice-cold 0.9% saline followed by freshly made 4% paraformaldehyde (PFA) in 0.1 M PBS. Brains were extracted, post-fixed for 24 hours at 4°C in 4% PFA and sunk in 30% sucrose in 0.1 M PBS over several days. Brains were stored at −80°C until they were ready to be sliced.

### 2.6. Histology and Immunohistochemistry

Frozen brains were cryosectioned at 40μm on an NX70 CryoStar Cryostat (Thermofischer Scientific). Free-floating sections were stored in 12 tubes with antifreeze, with every 12^th^ slice placed in the same tube. Samples were stored at −20°C prior to staining. Immunohistochemical (IHC) staining was conducted in order to quantify oligodendrocyte (OLs) and myelin markers. Specifically, IHC was used to detect myelin basic protein (MBP), one of the essential proteins in the myelin sheath (Hamano, Iwasaki, Takeya, & Takita, 1996) and glutathione s-transferase pi (GST-pi), a marker for immature to mature OLs (Tansey & Cammer, 1991). Tissue slices from one vial of tissue (every 12^th^ slice) were stained. Slices were first washed in tris-buffered saline (TBS), and blocked with 3% normal donkey serum (NDS) in TBS with 0.3% Triton-X100 for one hour at room temperature. Slices were then incubated overnight at 4°C with the following primary antibodies: rat anti-MBP (1:500, Abcam ab7349) and rabbit anti-GST-pi (1:5000, MBL 311). All antibodies were diluted in 0.3% TritonX-TBS containing 1% NDS. On day two, following three rinses in TBS (3 x 5 min), sections were incubated for two hours with the following secondary antibodies: AlexaFluor 488 donkey anti-rabbit and Cy3 donkey anti-rat (1:500, 711-545-152 and 712-165-153 respectively, Jackson ImmunoResearch Labs Inc). Sections were then rinsed in TBS (3 x 5 min) and incubated for 10 minutes with DAPI (1:40,000 in 1xPBS), followed by three more rinses in TBS. Lastly, sections were mounted on glass slides and cover slipped using 1,4 diazabicyclo[2.2.2]octane (DABCO).

### 2.7. Fluorescent Microscopy and Image Analysis

All tissue was imaged on an AxioScan Slide Scanner at 20x (AxioScan.Z1, Zeiss, Oberkochen, Germany). During analysis, experimenters were blind to animal condition. Regions of interest were chosen and exported as TIFFs using ZEN blue imaging software (Zeiss). For analysis of the prefrontal cortex, five PFC sections from AP 3.7 to AP 1.7 were used for analysis. Four regions of interest (ROIs) of the PFC were quantified: anterior cingulate cortex (ACC), prelimbic cortex (PL), infralimbic cortex (IL) and orbitofrontal cortex (OFC; Figure S1). For analysis of the amygdala, four sections from AP −2.00 to −3.24 were quantified, with ROIs for the basal amygdala (BA), lateral amygdala (LA) and central (CeA) amygdala (Figure S2). For the PFC and amygdala, all ROIs were hand-drawn in Fiji (Schindelin et al., 2012). For analysis of the hippocampus, six sections from AP −2.92 to −4.20 were used for analysis. Ten regions of the hippocampus were quantified: the CA1, CA2, CA3, lacunosum moleculare (LMol), molecular layer dentate gyrus (MoDG), oriens layer hippocampus (Or), radiatum layer hippocampus (Rad), stratum lucidum hippocampus (SLu), granule cell layer (GCL) and Hilus (Figure S3). The GCL and Hilus ROIs were hand-drawn in Fiji, however all of the other ROIs were analyzed by placing squares of varying sizes throughout the hippocampus. Squares were either 75 x 75 or 150 x 150 μm. All ROIs were placed based on consistent anatomical markers.

For all regions, GM myelin content was measured by using the integrated fluorescence intensity of MBP expression, normalized by area. OLs (GST-pi+ cells) were quantified using a custom written Fiji script, which included background subtraction, automated thresholding and particle analysis. The same parameters were used for all animals and all tissue regions. In addition, parameters were chosen such that the script counted within 10% of counts obtained by human experimenters. OL cell density is presented as the number of cells within a given ROI per mm^2^. In addition, cell density was calculated as a % of all DAPI+ cells; these data corresponded well with cell density by area, so all data presented are shown as cell density/area. Measures for each ROI were averaged across the anterior-posterior extent of sections and from both hemispheres.

### 2.8. Statistical Tests

All data are presented as the mean, plus or minus the standard error of the mean (SEM). Outliers were removed if they were more than two standard deviations above the group mean. For all 8 groups, across the 17 subregions and both markers (MBP and GST-pi), only 31 values were excluded. For analysis of corticosterone, we used a repeated measures ANOVA, followed by a Sidak’s post-hoc test for multiple comparisons. For analysis of GM myelin content and OLs, twoway ANOVA tests were used to compare male and female rats in the stress and control conditions at each age time point. An additional two-way ANOVA compared control vs. stress animals across different subregions of interest. Bonferroni post-hoc analyses were used to correct for multiple comparisons. Unpaired independent sample student’s t-tests were used to directly compare changes between control and stressed animals within each sex, when appropriate. Pearson correlations were used to identify relationships between corticosterone AUC and OL or myelin markers. In all tests, the alpha value was set at 0.05. All analyses were performed with GraphPad Prism version 8.4 (GraphPad Software, San Diego, California USA) and RStudio (RStudio Team, 2018).

## 3. Results

### 3.1. Male and female juvenile rats exhibit a robust physiological response following exposure to acute, severe stress

First, we examined the physiological response of male and female juvenile rats to acute stress exposure, in order to confirm our paradigm was indeed a potent stressor at this age. Previously, our lab and others have shown that predator scent coupled with immobilization produces a large corticosterone response in adult rats (Long et al., 2020; Morrow, Redmond, Roth, & Elsworth, 2000; Muroy, Long, Kaufer, & Kirby, 2016; Zoladz & Diamond, 2016). Here, exposure to three hours of immobilization stress in the presence of fox urine odor (Figure 1A) produced robust increases in serum corticosterone over baseline levels in both male and female juvenile rats (Figure 1B). The physiological increase in corticosterone observed 30 minutes into stress exposure persisted throughout the duration of the 3-hour stressor. A 2-way repeated measures ANOVA yielded a main effect of time (F (1.346, 40.39) = 90.49, p<0.0001), and a main effect of sex (F (1, 30) = 11.49, p<0.002), with no interaction between the two. A Sidak’s post-hoc test identified robust differences between corticosterone levels at baseline (0 min) compared to 30 and 180 minutes for both males and females (p<0.0001). Female rats demonstrated significantly higher levels of corticosterone relative to males at all time points, including at baseline (Figure 1B). This is consistent with previously reported findings that adult female rats exhibit higher baseline corticosterone levels (Kalil, Leite, Carvalho-Lima, & Anselmo-Franci, 2013; Mitsushima, Masuda, & Kimura, 2003). In addition to corticosterone, we assessed changes in body weight following acute stress exposure. Changes in body weight are strong physiological indicators of stress (Harris et al., 1998; Pulliam, Dawaghreh, Alema-Mensah, & Plotsky, 2010). Three days after stress, both male and female juveniles gained less weight relative to controls (males, t(28)=2.133, p=0.042; females, t(24)=2.246, p=0.034); Figure 1C). Collectively, these data suggest that acute, severe stress elicits a rapid physiological stress response in juvenile rats, increasing corticosterone levels and decreasing weight gain.

### 3.2. Myelin and oligodendrocytes are altered in limbic brain regions following acute juvenile stress in both the short and long term, with regional and sex specific differences

We sought to address whether acute severe juvenile stress affects grey matter (GM) myelin and oligodendrocytes (OLs) within three major limbic brain regions: the prefrontal cortex (PFC), amygdala (AMY) and hippocampus (HPC). In order to identify both short- and long-term effects of juvenile stress, one group of animals was sacrificed 12 days post stress exposure (at p40), while another was sacrificed almost two months post stress exposure, at an adult age (p95) (Figure 1A). We stained brain tissue for two markers: Glutathione S-transferase pi (GST-pi), a marker of immature to mature OLs, and myelin basic protein (MBP), a marker of myelination. In the PFC, we analyzed four separate subregions: the anterior cingulate cortex (ACC), the prelimbic (PL) and infralimbic (IL) cortices, and the orbitofrontal cortex (OFC) (Figure 1D). In the amygdala, we analyzed three subregions: the lateral amygdala (LA), basal amygdala (BA), and central amygdala (CeA) (Figure 1E). In the hippocampus, we analyzed ten subregions of interest: the CA1, CA2, CA3, lacunosum moleculare (LMol), molecular layer dentate gyrus (MoDG), oriens layer hippocampus (OR), radiatum layer hippocampus (RAD), stratum lucidum hippocampus (SLu), granule cell layer (GCL), and hilus regions (Figure 1F). Measures of OLs and GM myelin in the PFC, amygdala and hippocampus were averaged across subregions.

#### 3.2.1. Acute severe stress drives short-term increases in grey matter myelin in the amygdala and hippocampus of male rats

In all three brain regions (PFC, AMY and HPC), there were short term effects of stress on GM myelin, measured by MBP. In the PFC, there were no differences in MBP fluorescence intensity in males. For females, while we did not see an effect of stress within the whole PFC (p=0.08; Figure 2A), there was increased MBP, on average, relative to controls (Control = 25.8 ± 1.9 fluorescence/μm^2^, Stress = 30.8 ± 1.8 fluorescence/μ2) and a 2-way ANOVA (subregion x condition) revealed a main effect of stress (*F*(1,56)=11.04, p=0.0016; Table 1; Figure S4A). Stress-exposed female animals displayed higher levels of MBP in all subregions of the PFC compared to their respective controls, indicating the main effect of stress was not driven by any one PFC subregion (Table 1; Figure S4A). For both the amygdala and hippocampus, while there was no main effect of sex, there was an interaction between sex and condition, with increased MBP intensity in stress-exposed males and decreased MBP intensity in stress-exposed females (AMY: *F*(1,28)=10.78, p=0.0028; HPC: *F*(1,28) = 13.71, p=0.0009; Figures 2B-C). In the amygdala, MBP intensity significantly increased in stress-exposed males relative to controls (Males: Control = 37.2 ± 6.1 fluorescence/± μm^2^, Stress = 51.193 ± 7.5 fluorescence/μm^2^, *t*(14)=4.095, p=0.0011; Figure 2B). This effect of stress on MBP appeared across all of the amygdala ROIs. Bonferroni multiple comparisons tests revealed significant increases in MBP in the BA, LA, and CeA for stress-exposed males (BA: p=0.0097; LA: p=0.0104; CeA: p=0.0477; Figure S4B). In contrast to the males, stress-exposed females displayed, on average, reduced MBP; although this effect was not statistically significant averaged across the amygdala (p=0.10), there was a main effect of condition when testing across all amygdala subregions (*F*(1,42)=5.68, p=0.022; Table 1). In the hippocampus, there was a statistically significant increase in MBP in stress-exposed males compared to controls and a trend for decreased MBP relative to controls in stress-exposed females (Males: *t*(14)=4.745, p=0.000313; Females: *t*(14)=1.75, p=0.10; Figure 2C). This increased MBP in stress-exposed males occurred in the majority of individual subregions of the hippocampus, including in the CA1, CA2, CA3, MoDG, RAD, SLu and GCL (CA1: p = 0.0123; CA2: p = 0.0112; CA3: p = 0.0417; MoDG: p = 0.0066; RAD: p = 0.0228; SLu: p = 0.0142; GCL: p = 0.0307; Figure S5A). Interestingly, in stress-exposed females, there was a main effect of condition (*F*(1,138)=21.57, p<0.0001; Table 1) and multiple comparisons tests revealed significant decreases in MBP in the following hippocampal subregions: CA1, CA2, OR and RAD (CA1: p < 0.0001; CA2: p = 0.0470; OR: p = 0.0044; RAD: p = 0.0009; Figure S5B).

**Figure 2.**
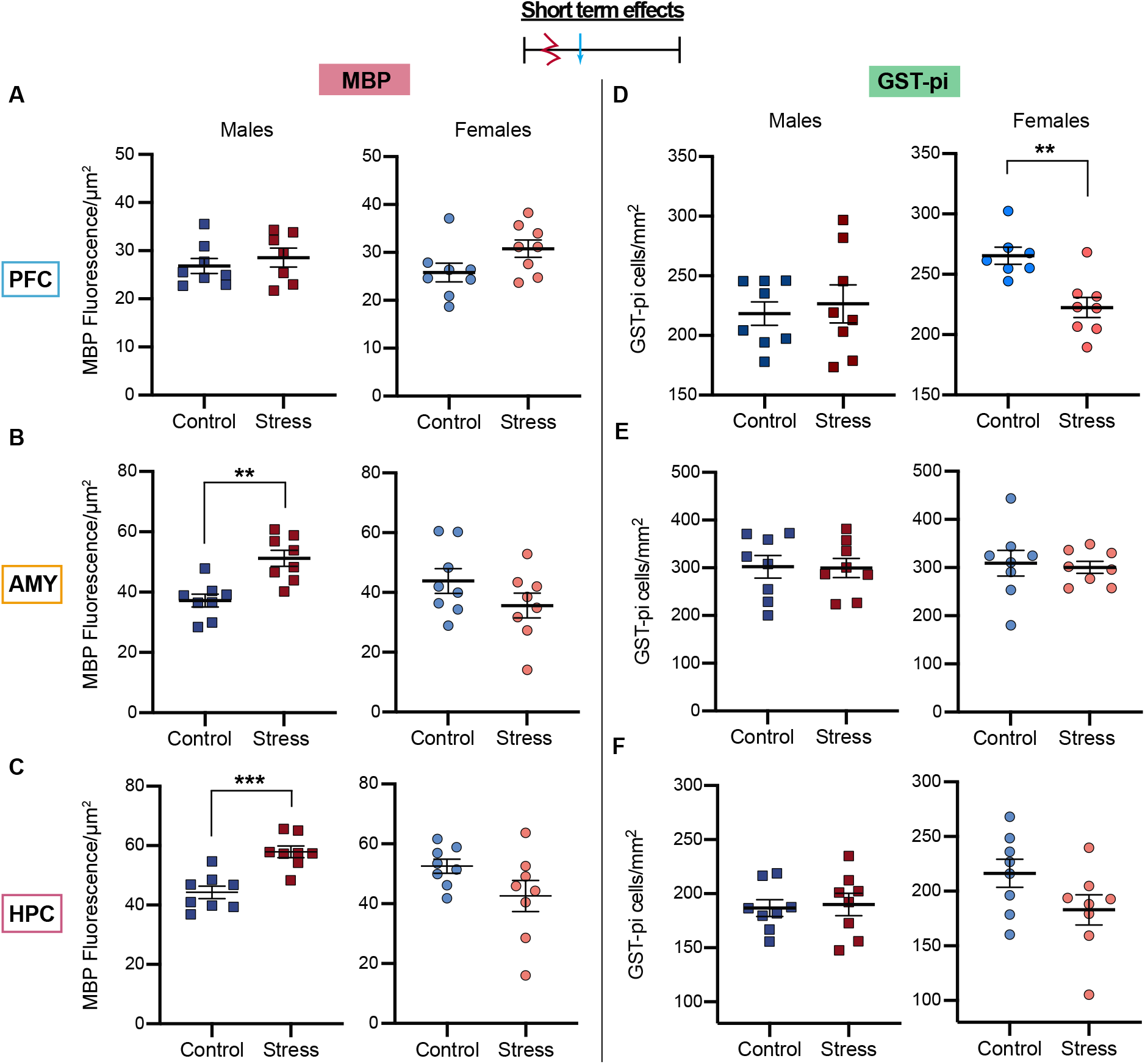
Short term effects of stress-exposure on myelin and oligodendrocytes. **A-C:** MBP fluorescence intensity results in the **(A)** PFC, **(B)** AMY and **(C)** HPC. **D-F:** GST-pi+ cell density results in the **(D)** PFC, **(E)** AMY and **(F)** HPC. Statistically significant differences are marked with asterisks (*p<0.05, ** p<0.01, *** p<0.001).

**Table 1.** Main effects of condition on MBP and GST-pi in limbic subregions of interest. Two-way ANOVAs (subregion x condition) were conducted for each major limbic region, testing for a main effect of condition (control or stress) on myelin and oligodendrocytes. F statistics and p-values are reported here. Statistically significant main effects are in bold.

|  | Short Term (p40) |  |  |  | Long term (p95) |  |  |  |
| --- | --- | --- | --- | --- | --- | --- | --- | --- |
|  | MBP |  | GST-pi |  | MBP |  | GST-pi |  |
|  | *Male* | *Female* | *Male* | *Female* | *Male* | *Female* | *Male* | *Female* |
| **PFC** | F(1,52)=1.49<br>p=0.23 | **F(1,56)=11.04**<br>**\*\*p=0.0016** | F(1,55)=0.77<br>p=0.38 | **F(1,52)=24.72**<br>**\*\*\*\*p<0.0001** | F(1,56)=0.52<br>p=0.47 | **F(1,56)=20.07**<br>**\*\*\*\*p<0.0001** | **F(1,56)=5.43**<br>**\*p=0.023** | F(1,54)=0.04<br>p=0.84 |
| **Amygdala** | **F(1,42)=43.23**<br>**\*\*\*\*p<0.0001** | **F(1,42)=5.68**<br>**\*p=0.022** | F(1,42)=0.01<br>p=0.92 | F(1,42)=0.16<br>p=0.69 | F(1,42)=0.05<br>p=0.83 | **F(1,42)=28.81**<br>**\*\*\*\*p<0.0001** | **F(1,41)=19.15**<br>**\*\*\*\*p<0.0001** | **F(1,41)=6.53**<br>**\*p=0.014** |
| **Hippocampus** | **F(1,138)=98.64**<br>**\*\*\*\*p<0.0001** | **F(1,138)=21.57**<br>**\*\*\*\*p<0.0001** | F(1,137)=0.35<br>p=0.56 | **F(1,135)=8.28**<br>**\*\*p=0.0047** | **F(1,140)=16.79**<br>**\*\*\*\*p<0.0001** | **F(1,139)=112.9**<br>**\*\*\*\*p<0.0001** | F(1,138)=3.20<br>p=0.08 | **F(1,136)=7.09**<br>**\*\*p=0.0087** |

#### 3.2.2. Acute severe stress drives short term decreases in oligodendrocytes in the PFC of female rats

Acute severe juvenile stress also produced short term changes in OL density across limbic regions in female rats. In the PFC, while there was no main effect of sex, there was an interaction between sex and condition (*F*(1,27)=5.3, p=0.0292). Stress-exposed females had reduced density of GST-pi+ cells in the PFC relative to controls (Control = 265 ± 7.1 cells/mm^2^, Stress = 222 ± 8.4 cells/mm^2^, *t*(13)=3.842, p=0.002, Figure 2D). No change was observed in stress-exposed males. Across the individual PFC subregions, there was again a main effect of stress for females (*F*(1,52)=24.72, p<0.0001; Table 1), but not males. A multiple comparisons test identified statistically significant decreases in GST-pi in the IL and OFC regions for stress-exposed females (p=0.0318 and p=0.0378 respectively; Figure S4C). In contrast, there were no changes in GST-pi density for males and females, both in the amygdala as a whole and in all amygdala subregions (Figure 2E, Figure S4D). Lastly, in the hippocampus, stress-exposed females tended to have decreased GST-pi+ density relative to controls (Control = 216 ± 36.205 cells/mm^2^, Stress = 182 ± 38.77 cells/mm^2^, p=0.09, Figure 2F) and indeed, there was a main effect of condition when looking across all hippocampal ROIs (*F*(1,135) = 8.28, p=0.0047; Table 1). Yet, there was also a large amount of variance in GST-pi across subregions (Figure S5). Decreased GST-pi density was seen across many subregions, and most robustly in the RAD (p=0.032). Yet other regions displayed no change or even increases relative to controls, such as in the CA2 and CA3 (Figure S5C-D). No changes were observed in males, either in the hippocampus as a whole or in any particular hippocampal ROIs.

Together, these findings suggest that acute severe stress leads to short term increases in GM myelin content, with increased myelin in the PFC in females and increased myelin in the amygdala and hippocampus for males. In contrast, OL density remained unchanged shortly following stress in all regions for males, while there were significant decreases in the hippocampus and PFC for females.

#### 3.2.3. Acute severe stress drives long term decreases in grey matter myelin across all limbic regions for female, but not male rats

We next looked at the long-term effects of acute severe stress exposure on GM myelin. At p95, two months after stress exposure, animals are well into adulthood. In all three brain regions, there were long term effects of juvenile stress on GM myelin. In particular, stress-exposed females, but not males, had reduced MBP fluorescence intensity in the PFC, AMY, and HPC compared to controls (PFC: *t*(11)=2.378, p=0.036, AMY: *t*(14)=3.362, p=0.0046, HPC: *t*(14)=4.262, p=0.0008, Figure 3A-C). In each of these areas, all subregions showed a female specific decrease in MBP intensity compared to control animals, highlighting main effects of stress (PFC: *F*(1,56) = 20.07, p<0.0001, AMY: *F*(1,42) = 28.81, p<0.0001, HPC: *F*(1,139) = 112.9, p<0.0001; Table 1). In the PFC, a Bonferroni multiple comparisons test identified significant decreases in MBP in the OFC region (p=0.022, Figure S6A). In the amygdala, all three ROIs showed significant decreases in MBP (BA: p=0.0014; LA: p=0.0070; CeA: p=0.0042; Figure S6B). Decreases in adult female hippocampal MBP also consistently appeared across multiple subregions of interest; all regions except the OR and SLu showed a significant decrease in MBP fluorescence intensity in stress compared to control animals (CA1: p=0.0005; CA2: p=0.0016; CA3: p=0.0002; LMol: p=0.0017; MoDG: p=0.0008; RAD: p=0.0137; GCL: p = 0.0010; Hilus: p<0.0001, Figure S7A-B). In contrast with females, no differences in average MBP were observed between stress-exposed and control adult males in any of three limbic regions (Figure 3A-C). Despite significant interactions between sex and condition in the PFC (*F*(1, 28)=4.4, p=0.0436) and the amygdala (*F*(1,28)=6.490, p=0.0166; Figure 3A-B), none of the PFC or amygdala subregions showed differences in MBP between stress exposed and control males. In hippocampal subregions however, there was a main effect of condition (*F*(1,140) = 16.79, p<0.0001; Table 1) and the OR showed a significant decrease in MBP levels in stress-exposed males (p=0.012; Figure S7A-B). Overall, juvenile acute stress produced robust long-term decreases in GM myelin in females, while males remained largely unaffected.

**Figure 3.**
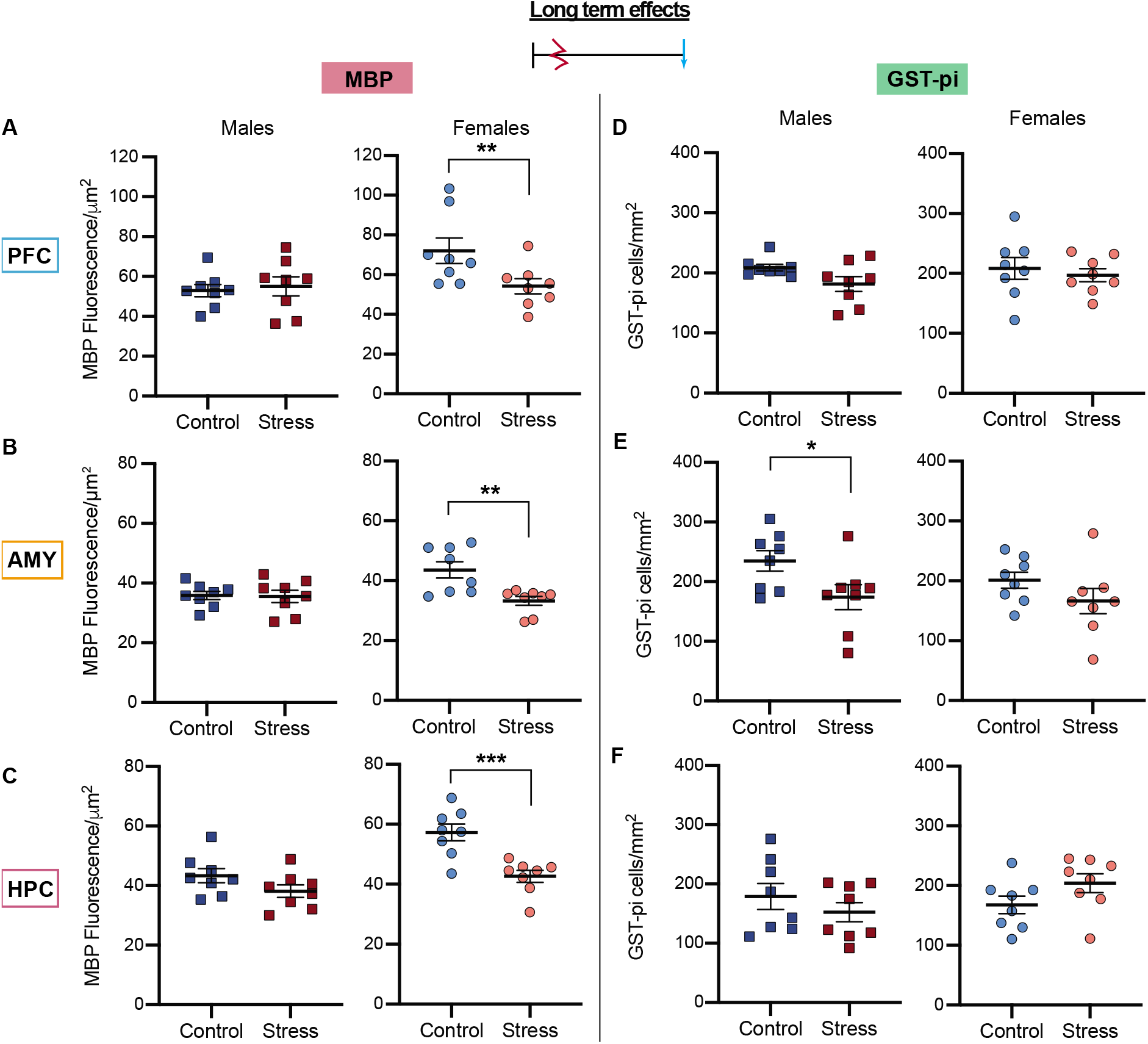
Long term effects of stress-exposure on myelin and oligodendrocytes. **A-C:** MBP fluorescence intensity results in the **(A)** PFC, **(B)** AMY and **(C)** HPC. **D-F:** GST-pi+ cell density results in the **(D)** PFC, **(E)** AMY and **(F)** HPC. Statistically significant differences are marked with asterisks (*p<0.05, ** p<0.01, ***p<0.001).

#### 3.2.4. Acute severe stress decreases oligodendrocyte density in the long term only in the CeA

Unlike effects identified in the short term, acute severe juvenile stress did not lead to consistent global changes in OL density across limbic regions. However, despite the lack of global effects, there were several significant changes in OL density at p95. In the PFC, stress-exposed males displayed a trend towards decreased GST-pi density (p=0.067; Figure 3D) that reflected a statistically significant main effect of stress on PFC subregions (*F*(1,56)=5.4, p=0.0234; Table 1). This mean decrease was observed for all subregions of the PFC (Figure S6C). In contrast, no changes in PFC GST-pi density were observed for adult females. In the amygdala, there was a significant decrease in GST-pi density for adult male rats (Male: *t*(14)=2.242, p=0.0417; Figure 3E), but not stress-exposed females. GST-pi density differed highly across subregions. Interestingly, GST-pi density was significantly lower in the CeA for both male and female stress-exposed animals relative to controls (Males: p=0.0059; Females: p=0.0118; Figure S6D). Lastly, in the hippocampus, no long-term change in average GST-pi density was observed when directly comparing controls to stress-exposed animals for either sex (Figure 3F). However, when looking at hippocampal subregions, there was a main effect of stress on GST-pi for females (*F*(1,136)=7.09, p=0.0087; Table 1), and on average, stress exposed females had a higher density of GST-pi+ cells relative to controls (Figure S7C-D).

Overall, for females, acute, severe stress as a juvenile leads to long term decreases in GM myelin content in limbic regions. In contrast, stress exposed males display no long-term changes in myelin levels. OL density is also largely unaffected, except for in the CeA, where both males and females display decreased OL density.

### 3.3. PFC, AMY and HPC myelin levels shortly after stress correlate with corticosterone responses

All juvenile animals exposed to acute stress demonstrated marked increases in corticosterone levels while undergoing the three-hour stressor, indicating a physiological stress response (Figure 1). Here, we explored whether these physiological changes in corticosterone corresponded with individual changes in GM myelin content and OL cell density. We assessed overall corticosterone levels utilizing the area under the curve (AUC). In the short term, at p40, AUC corticosterone was significantly correlated with myelin levels (MBP) in several groups (Figure 4). Specifically, in stress-exposed males, PFC myelin content was inversely correlated with corticosterone (r = −0.84, p=0.019; Figure 4A). Animals with the highest levels of corticosterone during stress-exposure had the lowest levels of PFC myelin at p40. In contrast, AMY and HPC myelin levels were positively correlated with AUC corticosterone in male animals. This effect was statistically significant in the amygdala and trending in the hippocampus (AMY: r = 0.89, p=0.003; HPC: r = 0.67, p=0.07; Figure 4B-C). In p40 stress-exposed females however, AUC corticosterone responses during stress-exposure were negatively correlated with myelin content in all three regions, though this relationship was only statistically significant in the hippocampus (r = −0.81, p=0.016; Figure 4C). For the AMY and HPC, these opposing correlational directions for males and females mimics the opposing directionality of group-wide changes in MBP; stress-exposed males had increased AMY and HPC myelin content at p40, while stress-exposed females had decreased AMY and HPC myelin content on average, relative to controls. Importantly, corticosterone levels at timepoint 0, which served as a baseline, were not significantly correlated with GM myelin content in p40 males or females for any of the brain regions analyzed. This indicates that corticosterone responses during stress exposure, and not basal corticosterone, was associated with subsequent myelin levels.

**Figure 4.**
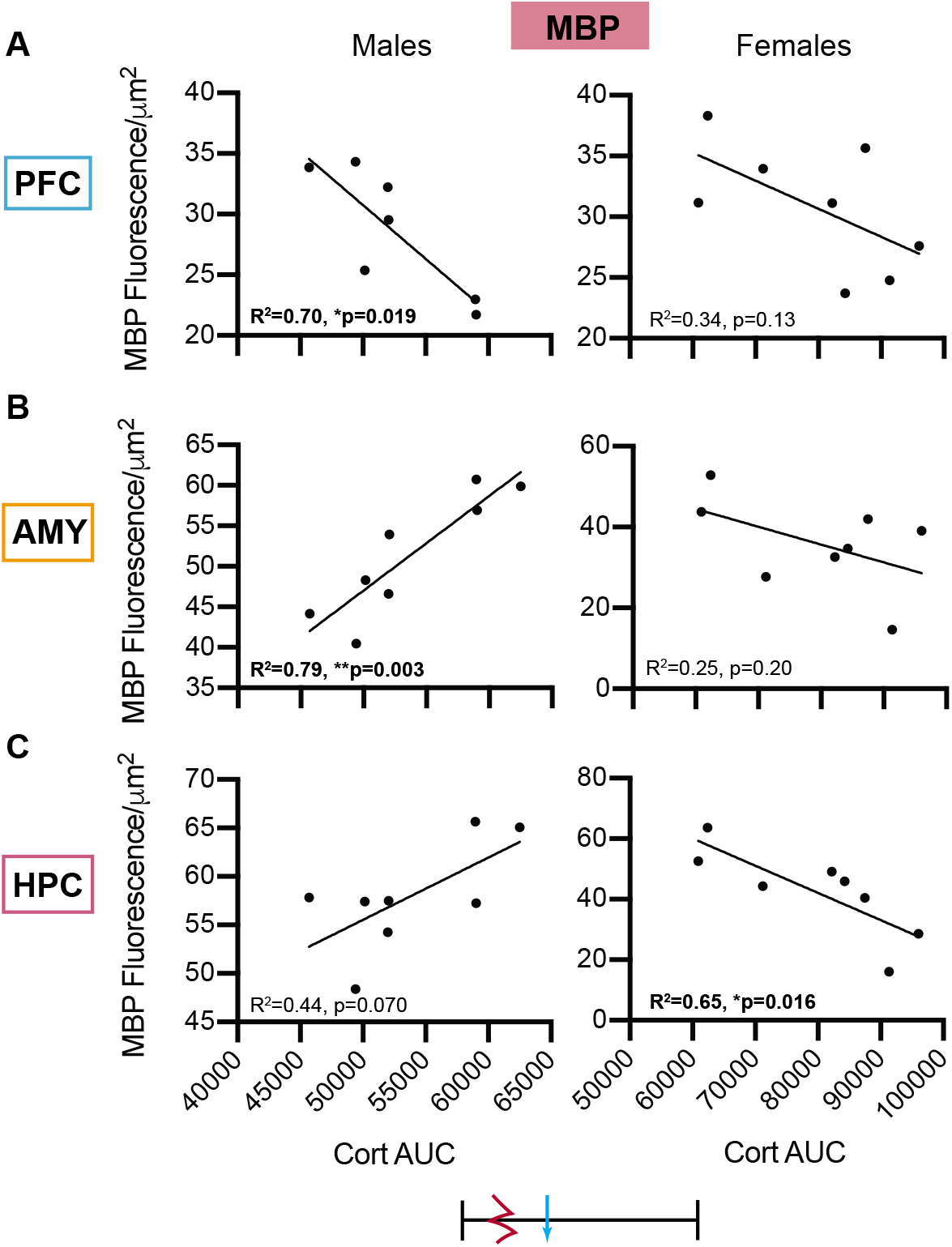
Correlations of corticosterone with short term myelin. Pearson’s correlations of MBP fluorescence intensity with area under the curve (AUC) for corticosterone in the **(A)** PFC, **(B)** AMY and **(C)** HPC of male and female rats tested at p40.

Although corticosterone was significantly associated with p40 MBP levels, there were no statistically significant correlations of corticosterone with GST-pi at p40, nor were there significant correlations of corticosterone with either MBP or GST-pi in the long term, in either sex (Figure S8). In addition, weight changes following stress did not strongly correlate with GM myelin content at either p40 or p95 (Figure S9). Overall, these data provide evidence that corticosterone levels during acute stress exposure may be associated with short-term PFC, AMY and HPC myelin levels.

## 4. Discussion

Changes in myelin and oligodendrocytes (OLs) are beginning to emerge as a novel mechanism that may contribute towards stress-induced pathologies (Chao et al., 2015; Chetty et al., 2014; Gibson, Geraghty, & Monje, 2018). Here, we aimed to fill a gap in the literature surrounding myelin and OL plasticity following juvenile trauma. In particular we focused on changes in the prefrontal cortex (PFC), amygdala (AMY) and hippocampus (HPC) - regions that are highly plastic and sensitive to stress during the peri-adolescent period (Spear, 2000). Furthermore, we sought to address if stress-induced alterations in grey matter (GM) myelin and OLs are long-lasting, and whether sex differences are observed. To our knowledge, this is the first study to examine changes in myelination in these regions after a single, acute stressor during this critical developmental window.

In this study, we exposed male and female juvenile rats to an acute traumatic stressor. First, we confirmed that juvenile stress exposure induced physiological changes; trauma increased serum corticosterone levels and reduced weight gain relative to controls in both males and females. Next, we tested whether exposure to acute severe stress as a juvenile produced subsequent short- or longterm changes in GM myelin and OLs in the PFC, AMY and HPC. Most interestingly, stress exposure altered myelin and OLs in all three brain regions, with changes occurring in a sex-specific manner. Further, exposure to juvenile stress had different effects on male and female myelin content and OLs across time. These changes are summarized in Table 2. In the short term, myelination in the PFC increased for stress exposed females, while AMY and HPC myelination increased for stress exposed males. Juvenile stress also reduced OL density in the PFC and HPC in p40 females but not males. In contrast to the short term increases in myelin, in the long term, juvenile stress decreased myelination in all three limbic regions exclusively in females. In the majority of brain regions, OL density remained largely unchanged in the long term. Altogether, our findings suggest that myelin and OLs in the PFC, AMY and HPC are sensitive to acute stress during the peri-adolescent time period of development. We conclude that experiencing a single acute severe stress as a juvenile alters myelin and OLs in a sex- and region-specific manner, with lasting impacts.

**Table 2.** Summary of stress-exposure effects on MBP and GST-pi across all conditions. Arrows denote direction of change. Bold arrows indicate statistically significant effects when comparing both the average of the region and the subregions of interest. Dashed arrows indicate effects that were trending when comparing the average of the region but showed a statistically significant main effect of condition when comparing the subregions of interest.

|  | Short Term (p40) |  |  |  | Long term (p95) |  |  |  |
| --- | --- | --- | --- | --- | --- | --- | --- | --- |
|  | MBP |  | GST-pi |  | MBP |  | GST-pi |  |
|  | *Male* | *Female* | *Male* | *Female* | *Male* | *Female* | *Male* | *Female* |
| **PFC** | -- | ↑ | -- | ↓ | -- | ↓ | ↓ | -- |
| **Amygdala** | ↑ | ↓ | -- | -- | -- | ↓ | ↓ | ↓ |
| **Hippocampus** | ↑ | ↓ | -- | ↓ | ↓ | ↓ | -- | ↑ |

### 4.1. Short-term changes following juvenile stress-exposure

In the short-term, stress-exposed females displayed increased levels of MBP across the PFC and males displayed increased MBP across the AMY and HPC. These findings supports the possibility that stress drives early maturation of limbic circuits (Bath, Manzano-Nieves, & Goodwill, 2016; Callaghan & Richardson, 2011; Gee et al., 2013; Honeycutt et al., 2020; Ono et al., 2008; Thomas, Caporale, Wu, & Wilbrecht, 2016). For example, in male mice, early life stress due to fragmented maternal care drives an earlier rise in MBP in the hippocampus (Bath et al., 2016), and early weaning prompts precocious myelination in the amygdala (Ono et al., 2008). During development, myelination acts to inhibit axonal sprouting and spine turnover, thereby acting as a brake on plasticity (Fields, 2008). Therefore, early myelination of the PFC, AMY or HPC could lead to impaired circuit functioning. In particular, plasticity within the PFC is especially critical for behaviors such as cognitive flexibility and decision making (Thomas et al., 2016). Alternatively, rather than driving early maturation to adult levels, stress may be altering developmental trajectories in the brain in a more transient manner (Thomas, Delevich, Chang, & Wilbrecht, 2020).

Our finding that myelination increases immediately following *acute* juvenile trauma stands in contrast with findings following exposure to *chronic* post-natal or juvenile stress. For example, chronic maternal separation reduced markers of myelin in the mPFC and hippocampus (Bordner et al., 2011; Wei et al., 2015; Yang et al., 2017). Furthermore, juvenile social isolation or social defeat reduced both myelin thickness and MBP levels in the mPFC (Liu et al., 2012; Zhang et al., 2016). Although GM myelination increased in several conditions shortly following exposure to acute juvenile stress, OL cell density either decreased or remained unchanged in the short term. In particular, OL density decreased in the PFC and HPC for stress exposed females, suggesting juvenile stress may impair OL proliferation or survival in those regions. A decrease in OL density following acute stress exposure is in line with reports of chronic early life adversity. For example, reductions in OLs are observed in the PFC of male mice following maternal separation (Bordner et al., 2011; Yang et al., 2017).

The most striking result in this study is that in females, but not in males, exposure as a juvenile to an acute traumatic event led to drastic and wide-spread alterations in the adult brain, measured as reductions in GM myelination across all three brain regions. The observed short-term decrease in OLs in females may partially explain this longer-term decrease in MBP; less immature and mature OLs at p40 might lead to fewer mature, myelinating OLs in adulthood as OLs undergo maturation over time (Noble, Mayer-Pröschel, & Miller, 2006). In our dataset, the same animals could not be examined at both time points due to the nature of our method; however, it would be interesting for future studies to analyze myelin in vivo in a longitudinal manner, perhaps with MRI imaging.

### 4.2. Long-term changes following juvenile stress-exposure

We observed a long-term reduction in PFC, AMY and HPC MBP in females, but no change in males. In prior work, adolescent social defeat did not change hippocampal myelin in the long term in male mice (Xu et al., 2020). In addition, in a study of adult stress in male mice, chronic social stress did not alter myelin related transcripts in the amygdala (Liu, Dietz, Hodes, Russo, & Casaccia, 2018). Thus, the lack of long-term changes in AMY and HPC myelin in male animals are not unexpected. Long term changes in AMY and HPC myelin in female animals, however, have never before been described, making our study the first to report a long lasting, acute stress-induced reduction in AMY and HPC myelination. A long-term reduction in PFC myelination is also in line with prior studies of juvenile chronic stress. For example, social isolation as a juvenile (from p21 to p35) led to reductions in PFC myelination, alterations in OL morphology and changes in mPFC-mediated behaviors when tested at p65 (Makinodan et al., 2012). Social isolation outside of this critical time window did not lead to such changes. We also found no changes in adult PFC OL cell number following juvenile stress. This is in line with a number of studies with stressors both in adolescence and adulthood that demonstrate reductions in PFC myelin levels but not OLs (Lehmann, Weigel, Elkahloun, & Herkenham, 2017; Liu et al., 2012; Makinodan et al., 2012; Xu et al., 2020; Zhang et al., 2016). A key difference, however, is that in these prior studies, changes in PFC myelination were observed in male, but not female animals, while here we observe the opposite effects. For example, one study found that social isolation (p30-p35) reduced MBP protein levels, quantified by western blots, in male but not female rats (Leussis & Andersen, 2008). Here, in contrast, in both the short and long-term, we found that the effects of traumatic stress on PFC myelination and OLs was limited to females. Different stressor types (social isolation or physical restraint stress), the timing of stress (whether before or after puberty onset), or the method of MBP assessment (immunohistochemistry or western blot), could all contribute to these observed dissimilarities. The majority of prior work focuses only on male rodents, and there remains a great need for side-by-side comparison of male and female animals using the same paradigms. Sex differences are especially important, as females are more likely to experience trauma during the peri-adolescent period, and trauma during this time has long-lasting consequences on mental health, such as increased risk for developing PTSD (Cuffe et al., 1998; Garza & Jovanovic, 2017; Udwin, Boyle, Yule, Bolton, & O’Ryan, 2000).

### 4.3. Associations of myelin with corticosterone responses during stress

Intriguingly, corticosterone responses during acute stress were correlated with myelin levels 12 days later at p40. The limbic system contains a high density of glucocorticoid receptors (GRs) and is highly sensitive to glucocorticoid signaling. OLs in these regions also express GRs (Chiba et al., 2012; Holsboer, 2000). Although a detailed mechanism remains unknown, there is literature to suggest glucocorticoids affect oligodendrogenesis and myelination in the adult brain (Chetty et al., 2014; Masters, Finch, & Nichols, 1994; Miyata et al., 2011; Wennström, Hellsten, Ekstrand, Lindgren, & Tingström, 2006). Corticosterone may therefore provide a possible mechanism by which stress alters myelination. In the PFC, for both males and females, animals with higher corticosterone responses showed the lowest levels of PFC myelination. Activation of GRs in the PFC may therefore either indirectly or directly contribute towards reduced myelination, although a causal role cannot be identified here. A different relationship was observed for AMY and HPC myelin. In males, higher corticosterone responses were associated with higher levels of AMY and HPC myelination. This is in line with prior work in our lab showing that corticosterone enhances oligodendrogenesis in the hippocampus of adult male rats (Chetty et al., 2014). It is possible that corticosterone may be driving increased myelination in these regions, leading to the group-wide short-term increase in AMY and HPC MBP in males. Yet, this correlation was reversed in female animals; animals with higher corticosterone showed reduced AMY and HPC myelin. This sex difference is interesting and merits further exploration in future studies. Females have a higher ratio of GRs to mineralcorticoid receptors (MRs), which could contribute towards this sex difference (Brydges et al., 2014). More broadly, males and females display different corticosterone responses following adolescent stress, perhaps helping explain some of the observed long-term sex differences. For example, adult female rats exposed to chronic stress during adolescence have higher basal corticosterone levels compared to nonstressed controls (Barha, Brummelte, Lieblich, & Galea, 2011). In contrast, chronic adolescent stress has no lasting impacts on basal corticosterone levels in adult male animals (Barha et al., 2011). This long-lasting change in basal corticosterone levels in females may be adaptive, or it may be suggestive of female-specific risk. Indeed, we observed here that adult females but not males, displayed reduced myelination across all limbic regions. Whether these changes are adaptive or not would need to be assessed through future work.

### 4.4. Limitations and Future Directions

While some limitations of the current study have already been discussed, there are several additional considerations. Although group effects were observed, the sample sizes were too low to reliably explore potential subgroups and inter-individual variations. Adding more timepoints would also improve resolution in the dataset, allowing us to map a curve of both GST-pi and MBP trajectories following stress, providing a deeper understanding into how stress alters myelin and OLs across time. In the current study we observed changes in MBP measured via fluorescence intensity. Increases in MBP intensity could reflect either an increase in the number of axons being myelinated, a thickening of already existing myelinated axons, or some combination of the two. Future work could use higher resolution imaging and/or electron microscopy to pull apart these possibilities and to address whether there are changes in OL morphology following acute juvenile stress. Higher resolution could also be applied to the local microcircuits within these limbic regions. For example, the PFC is a diverse brain region, with many subregions and local microcircuits across cortical layers (Dalley, Cardinal, & Robbins, 2004; Kolb, 1984). Although we looked at specific subregions of the PFC, each known to be involved in different aspects of behavior, we did not look at specific layers of the PFC. Prior literature has also focused on only deep layers of cortex (layers 5+6) where there are primarily pyramidal projection neurons (Lehmann et al., 2017; Makinodan et al., 2012). This is yet another difference between our study and others. Future work should aim to look at layer specific changes with a more detailed anatomical eye. Lastly, here, we looked at two specific markers: one for OLs and one for myelin. Future work could add additional markers of OLs to assess stress-induced changes across the OL lineage. OLs develop from oligodendrocyte progenitor cells (OPCs). OPCs persist throughout the postnatal period and continue to divide and generate myelinating OLs throughout their lifespan (Bergles & Richardson, 2016). While our study focused on immature and mature myelinating OLs, stress can also affect the progenitor pool in these limbic regions (Qiao et al., 2020; Saul et al., 2015; Teissier et al., 2020). For example, repeated variable stress selectively reduced the number of proliferating OPCs in the amygdala in juvenile male mice, while leaving proliferating neurons unchanged (Saul et al., 2015). Indeed, OPCs may play an important role in stress-related behaviors and should be analyzed in future work. For example, loss of OPCs in the PFC was sufficient to phenocopy depressive-like behaviors driven by chronic social stress (Birey et al., 2015).

OLs and myelin across limbic regions are now being recognized for their role in stress-associated behaviors. For example, adult PTSD patients have increased hippocampal myelin compared to trauma exposed controls (Chao et al., 2015). Interestingly, this increase in hippocampal myelin is positively correlated with PTSD symptom scores, suggesting that vulnerability to stress-induced disorders is related to hippocampal myelin. In a recent study from our lab, similar findings were observed in adult male rats. Specifically, OLs and GM myelin in the dentate gyrus of the hippocampus positively correlated with avoidance behaviors following exposure to acute severe stress. In addition, increased myelin levels in the amygdala were associated with enhanced fear behavior following trauma (Long et al., 2020). Together, these findings suggest that, in adults, OLs and myelin may be associated with individual vulnerability following acute stress. An important direction will be to assess whether changes in OLs and myelin following acute juvenile trauma correspond with subsequent changes in behavior. Acute severe stress in rodents also models another component of human PTSD; most animals will be resilient to stress, while a subset will display long lasting increases in fear and anxiety-like behaviors (Kessler et al., 1995; Russo, Murrough, Han, Charney, & Nestler, 2012; Zovkic, Meadows, Kaas, & Sweatt, 2013). Of particular interest then will be to see if changes in myelin and OLs are associated with susceptibility to juvenile acute stress. In a recent study, only animals susceptible to chronic social defeat had reduced myelin protein and fewer mature OLs in the mPFC. In addition, focal mPFC demyelination decreased social preference, implicating a causal role for myelin in social behavior (Bonnefil et al., 2019). Indeed, many prior studies measuring stress-induced changes in myelin and OLs have focused on associations with depressive-like behaviors, including social interaction (Birey et al., 2015; Lehmann et al., 2017; Leussis & Andersen, 2008; Liu et al., 2012, 2016; Tang et al., 2019). It will be interesting for future work to assess a diverse array of behaviors, covering the social, reward, fear and avoidance domains, to uncover how different behaviors affected by acute stress might be associated with myelin or OLs in a given limbic region.

The findings described here should also be considered within a broader circuit context. For example, the PFC, amygdala and hippocampus are all highly interconnected, and function together to regulate emotions (Ishikawa & Nakamura, 2003; Roozendaal et al., 2009). Connectivity between these regions is also altered following stress, in both humans and in rodents (Grandjean et al., 2016; Honeycutt et al., 2020). Myelination of axons corresponds with conduction velocity and synchronization across these brain regions (Monje, 2018; Pajevic & Basser, 2013). Thus, future studies should try to identify changes in myelination within a specific circuit, focusing on specific axonal projections between brain regions (for example, IL projections to the amygdala). Understanding how and if particular axonal projections are preferentially myelinated following stress will further our knowledge of circuit plasticity and its connection to behavior. A recent study found that pharmacological stimulation of neurons led to increases in myelination in an axon-specific manner. In addition, and relevant to the current study, juveniles showed higher sensitivity to stimulation than adults (Mitew et al., 2018). This suggests that neural activity, whether driven by stress or otherwise, may drive circuit specific modulation of myelin. Lastly, a critical area for future study will be to go beyond correlation and into causation, and to manipulate myelin and OLs directly within a brain region or circuit in order to assess their functional contribution to stress pathology and behaviors.

## 5. Conclusions

The findings presented here have important implications for understanding stress-sensitive developmental periods. There is increasing evidence that stress during late childhood and early adolescence may confer vulnerability for developing psychiatric disorders later in life (Carr et al., 2013; Heim & Nemeroff, 2001; Nemeroff, 2004a, 2004b; Ventriglio et al., 2015). Exposing rats to peri-adolescent stress is used to model the detrimental effects of childhood and early-adolescent trauma in humans (Tsoory, Cohen, & Richter-Levin, 2007; Tsoory & Richter-Levin, 2006). The majority of prior studies have tested chronic stressors during the peri-adolescent time period; however, many traumatic experiences are often acute in nature and can lead to long-lasting changes to the brain and behavior (Carrion & Wong, 2012; Nemeroff et al., 2006; Tsoory et al., 2007; Tsoory & Richter-Levin, 2006). Further, understanding acute trauma provides us with detailed knowledge about vulnerable windows during development when the brain is most sensitive to stress. Many studies have suggested that peri-adolescence is a sensitive period of development in which there is significant remodeling of limbic regions following stress (Spear, 2000). Our work here adds to the growing literature demonstrating that myelin and OLs are sensitive to stress early in life, providing an additional mechanism by which stress remodels the brain (Bath et al., 2016; Demaestri et al., 2020; Leussis & Andersen, 2008; Makinodan et al., 2012). Many psychiatric disorders, including schizophrenia, depression and PTSD, are characterized by alterations in myelination (Fields, 2008; Hamidi, Drevets, & Price, 2004; Lee & Fields, 2009; Lutz et al., 2017; Regenold et al., 2007; Tanti et al., 2018). However, whether changes in myelin contributes to vulnerability to these disorders, or whether they are simply a biomarker remains to be determined. Understanding stress-induced plasticity of PFC, AMY and HPC myelin and OLs may contribute to our understanding of these psychiatric disorders, as well as the vulnerability to developing pathology following early life stress. Studying sex differences in response to early life trauma is also important. In particular, females are known to have increased risk of developing stress-associated pathology, including PTSD (Gater et al., 1998; Kessler et al., 2005; Weissman et al., 2005). While behavior was not included in the current study, the selective long-term reductions in myelination in female animals is suggestive of female-specific risk. Overall, findings in rodents will inform our knowledge of how traumatic stressors may impact human brain development and mental health.

## Acknowledgments

We would like to thank Anjali Vadhri and Nora Galoustian for help with tissue processing and cell counting, and the many individuals who provided thoughtful edits, feedback and suggestions, including members of the Kaufer and Bath labs. Imaging was conducted at the CRL Molecular Imaging Center, supported by the UC Berkeley Biological Faculty Research Fund. We would like to thank Holly Aaron and Feather Ives for their microscopy training and assistance.

## Author contributions

Conceptualization: D.K. and J.B. Methodology: J.B., K.L., D.K. Investigation: J.B., M.B., K.H., S.F. Analysis: J.B., K.L., M.B., K.H., S.F. Visualizations & Writing: J.B., M.B., K.H., S.F., D.K. Resources, funding and supervision: D.K.

## Data and materials availability

All analyzed data associated with this study are available in the main text or in the supplementary materials. Raw data will be made available upon request.

## References

Andersen, S. L., Tomada, A., Vincow, E. S., Valente, E., Polcari, A., & Teicher, M. H. (2008). Preliminary Evidence for Sensitive Periods in the Effect of Childhood Sexual Abuse on Regional Brain Development. The Journal of Neuropsychiatry and Clinical Neurosciences, 20(3), 292–301. https://doi.org/10.1176/jnp.2008.20.3.292

Barha, C. K., Brummelte, S., Lieblich, S. E., & Galea, L. A. M. (2011). Chronic restraint stress in adolescence differentially influences hypothalamic-pituitary-adrenal axis function and adult hippocampal neurogenesis in male and female rats. Hippocampus. https://doi.org/10.1002/hipo.20829

Bath, K. G., Manzano-Nieves, G., & Goodwill, H. (2016). Early life stress accelerates behavioral and neural maturation of the hippocampus in male mice. Hormones and Behavior, 82(12), 64–71. https://doi.org/10.1016/j.yhbeh.2016.04.010

Bergles, D. E., & Richardson, W. D. (2016). Oligodendrocyte development and plasticity. Cold Spring Harbor Perspectives in Biology. https://doi.org/10.1101/cshperspect.a020453

Birey, F., Kloc, M., Chavali, M., Hussein, I., Wilson, M., Christoffel, D. J., … Aguirre, A. (2015). Genetic and Stress-Induced Loss of NG2 Glia Triggers Emergence of Depressive-like Behaviors through Reduced Secretion of FGF2. Neuron, 88(5), 941–956. https://doi.org/10.1016/j.neuron.2015.10.046

Birey, F., Kokkosis, A., & Aguirre, A. (2017). Oligodendroglia-lineage cells in brain plasticity, homeostasis and psychiatric disorders. Current Opinion in Neurobiology, 47, 93–103. https://doi.org/10.1016/j.conb.2017.09.016

Bolton, J. L., Short, A. K., Simeone, K. A., Daglian, J., & Baram, T. Z. (2019). Programming of stress-sensitive neurons and circuits by early-life experiences. Frontiers in Behavioral Neuroscience, 13(February), 1–9. https://doi.org/10.3389/fnbeh.2019.00030

Bonnefil, V., Dietz, K., Amatruda, M., Wentling, M., Aubry, A. V., Dupree, J. L., … Liu, J. (2019). Region-specific myelin differences define behavioral consequences of chronic social defeat stress in mice. ELife, 8, 1–13. https://doi.org/10.7554/eLife.40855

Bordner, K. A., George, E. D., Carlyle, B. C., Duque, A., Kitchen, R. R., Lam, T. K. T., … Simen, A. A. (2011). Functional genomic and proteomic analysis reveals disruption of myelin-related genes and translation in a mouse model of early life neglect. Frontiers in Psychiatry, 2(APR), 1–18. https://doi.org/10.3389/fpsyt.2011.00018

Breslau, N. (2009). The epidemiology of trauma, PTSD, and other posttrauma disorders. Trauma, Violence, and Abuse, 10(3), 198–210. https://doi.org/10.1177/1524838009334448

Brydges, N. M., Jin, R., Seckl, J., Holmes, M. C., Drake, A. J., & Hall, J. (2014). Juvenile stress enhances anxiety and alters corticosteroid receptor expression in adulthood. Brain and Behavior, 4(1), 4–13. https://doi.org/10.1002/brb3.182

Callaghan, B. L., & Richardson, R. (2011). Maternal separation results in early emergence of adult-like fear and extinction learning in infant rats. Behavioral Neuroscience. Callaghan, Bridget L.: School of Psychology, The University of New South Wales, Sydney, NSW, Australia, 2052,: American Psychological Association. https://doi.org/10.1037/a0022008

Carr, C. P., Martins, C. M. S., Stingel, A. M., Lemgruber, V. B., & Juruena, M. F. (2013). The role of early life stress in adult psychiatric disorders: A systematic review according to childhood trauma subtypes. Journal of Nervous and Mental Disease, 201(12), 1007–1020. https://doi.org/10.1097/NMD.0000000000000049

Carrion, V. G., Weems, C. F., & Reiss, A. L. (2007). Stress predicts brain changes in children: A pilot longitudinal study on youth stress, posttraumatic stress disorder, and the hippocampus. Pediatrics, 119(3), 509–516. https://doi.org/10.1542/peds.2006-2028

Carrion, V. G., & Wong, S. S. (2012). Can traumatic stress alter the brain? Understanding the implications of early trauma on brain development and learning. Journal of Adolescent Health, 51(2 SUPPL.), S23–S28. https://doi.org/10.1016/j.jadohealth.2012.04.010

Chao, L. L., Tosun, D., Woodward, S. H., Kaufer, D., & Neylan, T. C. (2015). Preliminary evidence of increased Hippocampal myelin content in veterans with posttraumatic stress disorder. Frontiers in Behavioral Neuroscience, 9(DEC), 1–8. https://doi.org/10.3389/fnbeh.2015.00333

Chetty, S., Friedman, A. R., Taravosh-Lahn, K., Kirby, E. D., Mirescu, C., Guo, F., … Kaufer, D. (2014). Stress and glucocorticoids promote oligodendrogenesis in the adult hippocampus. Molecular Psychiatry, 19(12), 1275–1283. https://doi.org/10.1038/mp.2013.190

Chiba, S., Numakawa, T., Ninomiya, M., Richards, M. C., Wakabayashi, C., & Kunugi, H. (2012). Chronic restraint stress causes anxiety-and depression-like behaviors, downregulates glucocorticoid receptor expression, and attenuates glutamate release induced by brain-derived neurotrophic factor in the prefrontal cortex. Progress in Neuro-Psychopharmacology and Biological Psychiatry, 39(1), 112–119. https://doi.org/ https://doi.org/10.1016/j.pnpbp.2012.05.018

Cohen, M. M., Jing, D., Yang, R. R., Tottenham, N., Lee, F. S., & Casey, B. J. (2013). Early-life stress has persistent effects on amygdala function and development in mice and humans. Proceedings of the National Academy of Sciences of the United States of America, 110(45), 18274–18278. https://doi.org/10.1073/pnas.1310163110

Cohen, R. A., Grieve, S., Hoth, K. F., Paul, R. H., Sweet, L., Tate, D., … Williams, L. M. (2006). Early Life Stress and Morphometry of the Adult Anterior Cingulate Cortex and Caudate Nuclei. Biological Psychiatry, 59(10), 975–982. https://doi.org/10.1016/j.biopsych.2005.12.016

Compas, B. E., & Phares, V. (1991). Stress during childhood and adolescence: Sources of risk and vulnerability. In Life-span developmental psychology: Perspectives on stress and coping. (pp. 111–129). Hillsdale, NJ, US: Lawrence Erlbaum Associates, Inc.

Cuffe, S. P., Addy, C. L., Garrison, C. Z., Waller, J. L., Jackson, K. L., McKeown, R. E., & Chilappagari, S. (1998). Prevalence of PTSD in a Community Sample of Older Adolescents. Journal of the American Academy of Child & Adolescent Psychiatry, 37(2), 147–154. https://doi.org/10.1097/00004583-199802000-00006

Dalley, J. W., Cardinal, R. N., & Robbins, T. W. (2004). Prefrontal executive and cognitive functions in rodents: Neural and neurochemical substrates. Neuroscience and Biobehavioral Reviews, 28(7), 771–784. https://doi.org/10.1016/j.neubiorev.2004.09.006

Demaestri, C., Pan, T., Critz, M., Ofray, D., Gallo, M., & Bath, K. G. (2020). Type of early life adversity confers differential, sex-dependent effects on early maturational milestones in mice. Hormones and Behavior, 124(February), 104763. https://doi.org/10.1016/j.yhbeh.2020.104763

Eiland, L., Ramroop, J., Hill, M. N., Manley, J., & McEwen, B. S. (2012). Chronic juvenile stress produces corticolimbic dendritic architectural remodeling and modulates emotional behavior in male and female rats. Psychoneuroendocrinology, 37(1), 39–47. https://doi.org/ https://doi.org/10.1016/j.psyneuen.2011.04.015

Eiland, L., & Romeo, R. D. (2013). Stress and the developing adolescent brain. Neuroscience, 249, 162–171. https://doi.org/10.1016/j.neuroscience.2012.10.048

Ellis, B. J., & Boyce, W. T. (2008). Biological sensitivity to context. Current Directions in Psychological Science, 17(3), 183–187. https://doi.org/10.1111/j.1467-8721.2008.00571.x

Falkai, P., Malchow, B., Wetzestein, K., Nowastowski, V., Bernstein, H.-G., Steiner, J., … Schmitt, A. (2016). Decreased Oligodendrocyte and Neuron Number in Anterior Hippocampal Areas and the Entire Hippocampus in Schizophrenia: A Stereological Postmortem Study. Schizophrenia Bulletin, 42(suppl_1), S4–S12. https://doi.org/10.1093/schbul/sbv157

Fields, R. D. (2008). White matter in learning, cognition and psychiatric disorders. Trends in Neurosciences, 31(7), 361–370. https://doi.org/10.1016/j.tins.2008.04.001

Garza, K., & Jovanovic, T. (2017). Impact of Gender on Child and Adolescent PTSD. Current Psychiatry Reports, 19(11), 87. https://doi.org/10.1007/s11920-017-0830-6

Gater, R., Tansella, M., Korten, A., Tiemens, B. G., Mavreas, V. G., & Olatawura, M. O. (1998). Sex Differences in the Prevalence and Detection of Depressive and Anxiety Disorders in General Health Care Settings: Report From the World Health Organization Collaborative Study on Psychological Problems in General Health Care. Archives of General Psychiatry, 55(5), 405–413. https://doi.org/10.1001/archpsyc.55.5.405

Gee, D. G., & Casey, B. J. (2015). The impact of developmental timing for stress and recovery. Neurobiology of Stress, 1, 184–194. https://doi.org/ https://doi.org/10.1016/j.ynstr.2015.02.001

Gee, D. G., Gabard-Durnam, L. J., Flannery, J., Goff, B., Humphreys, K. L., Telzer, E. H., … Tottenham, N. (2013). Early developmental emergence of human amygdala–prefrontal connectivity after maternal deprivation. Proceedings of the National Academy of Sciences, 110(39), 15638 LP – 15643. https://doi.org/10.1073/pnas.1307893110

Gibson, E. M., Geraghty, A. C., & Monje, M. (2018). Bad wrap: Myelin and myelin plasticity in health and disease. Developmental Neurobiology, 78(2), 123–135. https://doi.org/10.1002/dneu.22541

Grandjean, J., Azzinnari, D., Seuwen, A., Sigrist, H., Seifritz, E., Pryce, C. R., & Rudin, M. (2016). Chronic psychosocial stress in mice leads to changes in brain functional connectivity and metabolite levels comparable to human depression. NeuroImage. https://doi.org/10.1016/j.neuroimage.2016.08.013

Gutman, D. A., & Nemeroff, C. B. (2002). Neurobiology of early life stress: rodent studies. Seminars in Clinical Neuropsychiatry, 7(2), 89–95. https://doi.org/10.1053/scnp.2002.31781

Hamano, K., Iwasaki, N., Takeya, T., & Takita, H. (1996). A quantitative analysis of rat central nervous system myelination using the immunohistochemical method for MBP. Developmental Brain Research, 93(1–2), 18–22. https://doi.org/10.1016/0165-3806(96)00025-9

Hamidi, M., Drevets, W. C., & Price, J. L. (2004). Glial reduction in amygdala in major depressive disorder is due to oligodendrocytes. Biological Psychiatry, 55(6), 563–569. https://doi.org/10.1016/j.biopsych.2003.11.006

Hanson, J. L., Adluru, N., Chung, M. K., Alexander, A. L., Davidson, R. J., & Pollak, S. D. (2013). Early neglect is associated with alterations in white matter integrity and cognitive functioning. Child Development, 84(5), 1566–1578. https://doi.org/10.1111/cdev.12069

Harris, R. B. S., Zhou, J., Youngblood, B. D., Rybkin, I. I., Smagin, G. N., & Ryan, D. H. (1998). Effect of repeated stress on body weight and body composition of rats fed low-and high-fat diets. American Journal of Physiology - Regulatory Integrative and Comparative Physiology, 275(6 44-6). https://doi.org/10.1152/ajpregu.1998.275.6.r1928

Heim, C., & Nemeroff, C. B. (2001). The role of childhood trauma in the neurobiology of mood and anxiety disorders: Preclinical and clinical studies. Biological Psychiatry, 49(12), 1023–1039. https://doi.org/10.1016/S0006-3223(01)01157-X

Holsboer, F. (2000). The corticosteroid receptor hypothesis of depression. Neuropsychopharmacology. https://doi.org/10.1016/S0893-133X(00)00159-7

Honeycutt, J. A., Demaestri, C., Peterzell, S., Silveri, M. M., Cai, X., Kulkarni, P., … Brenhouse, H. C. (2020). Altered corticolimbic connectivity reveals sex-specific adolescent outcomes in a rat model of early life adversity. ELife, 9, 1–27. https://doi.org/10.7554/eLife.52651

Hughes, K., Bellis, M. A., Hardcastle, K. A., Sethi, D., Butchart, A., Mikton, C., … Dunne, M. P. (2017). The effect of multiple adverse childhood experiences on health: a systematic review and meta-analysis. The Lancet Public Health, 2(8), e356–e366. https://doi.org/ https://doi.org/10.1016/S2468-2667(17)30118-4

Isgor, C., Kabbaj, M., Akil, H., & Watson, S. J. (2004). Delayed effects of chronic variable stress during peripubertal-juvenile period on hippocampal morphology and on cognitive and stress axis functions in rats. Hippocampus, 14(5), 636–648. https://doi.org/10.1002/hipo.10207

Ishikawa, A., & Nakamura, S. (2003). Convergence and Interaction of Hippocampal and Amygdalar Projections within the Prefrontal Cortex in the Rat. Journal of Neuroscience, 23(31), 9987–9995. https://doi.org/10.1523/jneurosci.23-31-09987.2003

Johnson, F. K., Delpech, J. C., Thompson, G. J., Wei, L., Hao, J., Herman, P., … Kaffman, A. (2018). Amygdala hyper-connectivity in a mouse model of unpredictable early life stress. Translational Psychiatry, 8(1). https://doi.org/10.1038/s41398-018-0092-z

Kalil, B., Leite, C. M., Carvalho-Lima, M., & Anselmo-Franci, J. A. (2013). Role of sex steroids in progesterone and corticosterone response to acute restraint stress in rats: Sex differences. Stress, 16(4), 452–460. https://doi.org/10.3109/10253890.2013.777832

Kessler, R. C., Amminger, G. P., Aguilar-Gaxiola, S., Alonso, J., Lee, S., & Üstün, T. B. (2007). Age of onset of mental disorders: A review of recent literature. Current Opinion in Psychiatry, 20(4), 359–364. https://doi.org/10.1097/YCO.0b013e32816ebc8c

Kessler, R. C., Berglund, P., Demler, O., Jin, R., Merikangas, K. R., & Walters, E. E. (2005). Lifetime Prevalence and Age-of-Onset Distributions of. Arch Gen Psychiatry, 62(June), 593–602. https://doi.org/10.1001/archpsyc.62.6.593

Kessler, R. C., Sonnega, A., Bromet, E., Hughes, M., & Nelson, C. B. (1995). Posttraumatic Stress Disorder in the National Comorbidity Survey. Archives of General Psychiatry, 52(12), 1048–1060. https://doi.org/10.1001/archpsyc.1995.03950240066012

Kolb, B. (1984). Functions of the frontal cortex of the rat: a comparative review. Brain Research, 320(1), 65–98.

Lee, P., & Fields, D. (2009). Regulation of myelin genes implicated in psychiatric disorders by functional activity in axons. Frontiers in Neuroanatomy. Retrieved from https://www.frontiersin.org/article/10.3389/neuro.05.004.2009

Lehmann, M. L., Weigel, T. K., Elkahloun, A. G., & Herkenham, M. (2017). Chronic social defeat reduces myelination in the mouse medial prefrontal cortex. Scientific Reports, 7(March), 1–13. https://doi.org/10.1038/srep46548

Leussis, M. P., & Andersen, S. L. (2008). Is adolescence a sensitive period for depression? Behavioral and neuroanatomical findings from a social stress model. Synapse, 62(1), 22–30. https://doi.org/10.1002/syn.20462

Liu, J., Dietz, K., DeLoyht, J. M., Pedre, X., Kelkar, D., Kaur, J., … Casaccia, P. (2012). Impaired adult myelination in the prefrontal cortex of socially isolated mice. Nature Neuroscience, 15(12), 1621–1623. https://doi.org/10.1038/nn.3263

Liu, J., Dietz, K., Hodes, G. E., Russo, S. J., & Casaccia, P. (2018). Widespread transcriptional alternations in oligodendrocytes in the adult mouse brain following chronic stress. Developmental Neurobiology, 78(2), 152–162. https://doi.org/10.1002/dneu.22533

Liu, J., Dupree, J. L., Gacias, M., Frawley, R., Sikder, T., Naik, P., & Casaccia, P. (2016). Clemastine enhances myelination in the prefrontal cortex and rescues behavioral changes in socially isolated mice. Journal of Neuroscience, 36(3), 957–962. https://doi.org/10.1523/JNEUROSCI.3608-15.2016

Long, K., Breton, J., Chao, L., Sorooshyari, S., Hu, K., An, A., … Kaufer, D. (2020). Region-Specific Maladaptive Myelination Contributes to Differential Susceptibility to Stress-Induced Avoidance and Acute Threat Reactivity in Humans and Rodents. Biological Psychiatry, 87, S88. https://doi.org/10.1016/j.biopsych.2020.02.247

Luby, J., Belden, A., Botteron, K., Marrus, N., Harms, M. P., Babb, C., … Barch, D. (2013). The effects of poverty on childhood brain development: The mediating effect of caregiving and stressful life events. JAMA Pediatrics, 167(12), 1135–1142. https://doi.org/10.1001/jamapediatrics.2013.3139

Lutz, P. E., Tanti, A., Gasecka, A., Barnett-Burns, S., Kim, J. J., Zhou, Y., … Turecki, G. (2017). Association of a history of child abuse with impaired myelination in the anterior cingulate cortex: Convergent epigenetic, transcriptional, and morphological evidence. American Journal of Psychiatry, 174(12), 1185–1194. https://doi.org/10.1176/appi.ajp.2017.16111286

Ma, N., Li, L., Shu, N., Liu, J., Gong, G., He, Z., … Jiang, T. (2007). White Matter Abnormalities in First-Episode, Treatment-Naive Young Adults With Major Depressive Disorder. American Journal of Psychiatry, 164(5), 823–826. https://doi.org/10.1176/ajp.2007.164.5.823

Makinodan, M., Rosen, K. M., Ito, S., & Corfas, G. (2012). A Critical Period for Social Experience-Dependent Oligodendrocyte Maturation and Myelination. Science, 337(6100), 1357–1360. https://doi.org/10.1126/science.1220845

Masters, J. N., Finch, C. E., & Nichols, N. R. (1994). Rapid Increase in Glycerol Phosphate Dehydrogenase mRNA in Adult Rat Brain: A Glucocorticoid-Dependent Stress Response. Neuroendocrinology, 60(1), 23–35. https://doi.org/10.1159/000126716

McCrory, E. J., De Brito, S. A., Kelly, P. A., Bird, G., Sebastian, C. L., Mechelli, A., … Viding, E. (2013). Amygdala activation in maltreated children during pre-attentive emotional processing. British Journal of Psychiatry, 202(4), 269–276. https://doi.org/DOI:10.1192/bjp.bp.112.116624

McEwen, B. S., Bowles, N. P., Gray, J. D., Hill, M. N., Hunter, R. G., Karatsoreos, I. N., & Nasca, C. (2015). Mechanisms of stress in the brain. Nature Neuroscience, 18(10), 1353–1363. https://doi.org/10.1038/nn.4086

McEwen, B. S., & Stellar, E. (1993). Stress and the Individual: Mechanisms Leading to Disease. Archives of Internal Medicine, 153(18), 2093–2101. https://doi.org/10.1001/archinte.1993.00410180039004

McLean, C. P., & Anderson, E. R. (2009). Brave men and timid women? A review of the gender differences in fear and anxiety. Clinical Psychology Review, 29(6), 496–505. https://doi.org/10.1016/j.cpr.2009.05.003

Meaney, M. J., Aitken, D. H., Van Berkel, C., Bhatnagar, S., & Sapolsky, R. M. (1988). Effect of neonatal handling on age-related impairments associated with the hippocampus. Science, 239(4841), 766–768. https://doi.org/10.1126/science.3340858

Mitew, S., Gobius, I., Fenlon, L. R., McDougall, S. J., Hawkes, D., Xing, Y. L., … Emery, B. (2018). Pharmacogenetic stimulation of neuronal activity increases myelination in an axon-specific manner. Nature Communications, 9(1), 1–16. https://doi.org/10.1038/s41467-017-02719-2

Mitsushima, D., Masuda, J., & Kimura, F. (2003). Sex differences in the stress-induced release of acetylcholine in the hippocampus and corticosterone from the adrenal cortex in rats. Neuroendocrinology, 78(4), 234–240. https://doi.org/10.1159/000073707

Miyata, S., Koyama, Y., Takemoto, K., Yoshikawa, K., Ishikawa, T., Taniguchi, M., … Tohyama, M. (2011). Plasma corticosterone activates SGK1 and induces morphological changes in oligodendrocytes in corpus callosum. PLoS ONE, 6(5). https://doi.org/10.1371/journal.pone.0019859

Monje, M. (2018). Myelin Plasticity and Nervous System Function. Annual Review of Neuroscience, 41(1), 61–76. https://doi.org/10.1146/annurev-neuro-080317-061853

Morrow, B. A., Redmond, A. J., Roth, R. H., & Elsworth, J. D. (2000). The predator odor, TMT, displays a unique, stress-like pattern of dopaminergic and endocrinological activation in the rat. Brain Research. https://doi.org/10.1016/S0006-8993(00)02174-0

Muroy, S. E., Long, K. L. P., Kaufer, D., & Kirby, E. D. (2016). Moderate Stress-Induced Social Bonding and Oxytocin Signaling are Disrupted by Predator Odor in Male Rats. Neuropsychopharmacology, 41(8), 2160–2170. https://doi.org/10.1038/npp.2016.16

Nave, K. A., & Ehrenreich, H. (2014). Myelination and oligodendrocyte functions in psychiatric diseases. JAMA Psychiatry, 71(5), 582–584. https://doi.org/10.1001/jamapsychiatry.2014.189

Nemeroff, C. B. (2004a). Early-Life Adversity, CRF Dysregulation, and Vulnerability to Mood and Anxiety Disorders. Psychopharmacology Bulletin, 38(1), 14–20. Retrieved from http://europepmc.org/abstract/MED/15278013

Nemeroff, C. B. (2004b). Neurobiological consequences of childhood trauma. The Journal of Clinical Psychiatry. Nemeroff, Charles B.: Emory University School of Medicine, 1639 Pierce Dr., Ste. 4000, Atlanta, GA, US, 30322–4990,: Physicians Postgraduate Press.

Nemeroff, C. B., Bremner, J. D., Foa, E. B., Mayberg, H. S., North, C. S., & Stein, M. B. (2006). Posttraumatic stress disorder: A state-of-the-science review. Journal of Psychiatric Research, 40(1), 1–21. https://doi.org/10.1016/j.jpsychires.2005.07.005

Noble, M., Mayer-Pröschel, M., & Miller, R. H. (2006). The Oligodendrocyte. Developmental Neurobiology, 151–196. https://doi.org/10.1007/0-387-28117-7_6

Nooner, K. B., Mennes, M., Brown, S., Castellanos, F. X., Leventhal, B., Milham, M. P., & Colcombe, S. J. (2013). Relationship of Trauma Symptoms to Amygdala-Based Functional Brain Changes in Adolescents. Journal of Traumatic Stress, 26(6), 784–787. https://doi.org/10.1002/jts.21873

Ono, M., Kikusui, T., Sasaki, N., Ichikawa, M., Mori, Y., & Murakami-Murofushi, K. (2008). Early weaning induces anxiety and precocious myelination in the anterior part of the basolateral amygdala of male Balb/c mice. Neuroscience, 156(4), 1103–1110. https://doi.org/10.1016/j.neuroscience.2008.07.078

Oztan, O., Aydin, C., & Isgor, C. (2011). Chronic variable physical stress during the peripubertal-juvenile period causes differential depressive and anxiogenic effects in the novelty-seeking phenotype: functional implications for hippocampal and amygdalar brain-derived neurotrophic factor and th. Neuroscience, 192(1), 334–344. https://doi.org/10.1016/j.neuroscience.2011.06.077

Pagliaccio, D., Luby, J. L., Bogdan, R., Agrawal, A., Gaffrey, M. S., Belden, A. C., … Barch, D. M. (2014). Stress-system genes and life stress predict cortisol levels and amygdala and hippocampal volumes in children. Neuropsychopharmacology, 39(5), 1245–1253. https://doi.org/10.1038/npp.2013.327

Pajevic, S., & Basser, P. J. (2013). An Optimum Principle Predicts the Distribution of Axon Diameters in Normal White Matter. PLOS ONE, 8(1), e54095. Retrieved from https://doi.org/10.1371/journal.pone.0054095

Piekarski, D. J., Johnson, C. M., Boivin, J. R., Thomas, A. W., Lin, W. C., Delevich, K., … Wilbrecht, L. (2017). Does puberty mark a transition in sensitive periods for plasticity in the associative neocortex? Brain Research, 1654, 123–144. https://doi.org/10.1016/j.brainres.2016.08.042

Popoli, M., Yan, Z., McEwen, B. S., & Sanacora, G. (2012). The stressed synapse: the impact of stress and glucocorticoids on glutamate transmission. Nature Reviews Neuroscience, 13(1), 22–37. https://doi.org/10.1038/nrn3138

Pulliam, J. V. K., Dawaghreh, A. M., Alema-Mensah, E., & Plotsky, P. M. (2010). Social defeat stress produces prolonged alterations in acoustic startle and body weight gain in male Long Evans rats. Journal of Psychiatric Research, 44(2), 106–111. https://doi.org/10.1016/j.jpsychires.2009.05.005

Qiao, Y., Zhao, J., Li, C., Zhang, M., Wei, L., Zhang, X., … Gao, T. (2020). Effect of combined chronic predictable and unpredictable stress on depression-like symptoms in mice. Annals of Translational Medicine. https://doi.org/10.21037/atm-20-5168

Regenold, W. T., Phatak, P., Marano, C. M., Gearhart, L., Viens, C. H., & Hisley, K. C. (2007). Myelin staining of deep white matter in the dorsolateral prefrontal cortex in schizophrenia, bipolar disorder, and unipolar major depression. Psychiatry Research, 151(3), 179–188. https://doi.org/10.1016/j.psychres.2006.12.019

Roozendaal, B., McEwen, B. S., & Chattarji, S. (2009). Stress, memory and the amygdala. Nature Reviews Neuroscience. https://doi.org/10.1038/nrn2651

RStudio Team,. (2018). RStudio: Integrated Development for R. Boston, MA: PBC. Retrieved from http://www.rstudio.com/

Russo, S. J., Murrough, J. W., Han, M. H., Charney, D. S., & Nestler, E. J. (2012). Neurobiology of resilience. Nature Neuroscience, 15(11), 1475–1484. https://doi.org/10.1038/nn.3234

Saul, M. L., Helmreich, D. L., Rehman, S., & Fudge, J. L. (2015). Proliferating cells in the adolescent rat amygdala: Characterization and response to stress. Neuroscience, 311, 105–117. https://doi.org/ https://doi.org/10.1016/j.neuroscience.2015.10.003

Schindelin, J., Arganda-Carreras, I., Frise, E., Kaynig, V., Longair, M., Pietzsch, T., … Cardona, A. (2012). Fiji: An open-source platform for biological-image analysis. Nature Methods, 9(7), 676–682. https://doi.org/10.1038/nmeth.2019

Sokolov, B. P. (2007). Oligodendroglial abnormalities in schizophrenia, mood disorders and substance abuse. Comorbidity, shared traits, or molecular phenocopies? International Journal of Neuropsychopharmacology, 10(4), 547–555. https://doi.org/10.1017/S1461145706007322

Spear, L. P. (2000). The adolescent brain and age-related behavioral manifestations. Neuroscience & Biobehavioral Reviews, 24(4), 417–463. https://doi.org/ https://doi.org/10.1016/S0149-7634(00)00014-2

Tang, J., Liang, X., Zhang, Y., Chen, L., Wang, F., Tan, C., … Tang, Y. (2019). The effects of running exercise on oligodendrocytes in the hippocampus of rats with depression induced by chronic unpredictable stress. Brain Research Bulletin, 149(November 2018), 1–10. https://doi.org/10.1016/j.brainresbull.2019.04.001

Tansey, F. A., & Cammer, W. (1991). A Pi Form of Glutathione-S-Transferase Is a Myelin-and Oligodendrocyte-Associated Enzyme in Mouse Brain. Journal of Neurochemistry, 57(1), 95–102. https://doi.org/10.1111/j.1471-4159.1991.tb02104.x

Tanti, A., Kim, J. J., Wakid, M., Davoli, M. A., Turecki, G., & Mechawar, N. (2018). Child abuse associates with an imbalance of oligodendrocyte-lineage cells in ventromedial prefrontal white matter. Molecular Psychiatry, 23(10), 2018–2028. https://doi.org/10.1038/mp.2017.231

Teicher, M. H., Samson, J. A., Polcari, A., & McGreenery, C. E. (2006). Sticks, stones, and hurtful words: Relative effects of various forms of childhood maltreatment. American Journal of Psychiatry, 163(6), 993–1000. https://doi.org/10.1176/ajp.2006.163.6.993

Teissier, A., Le Magueresse, C., Olusakin, J., Andrade da Costa, B. L. S., De Stasi, A. M., Bacci, A., … Gaspar, P. (2020). Early-life stress impairs postnatal oligodendrogenesis and adult emotional behaviour through activity-dependent mechanisms. Molecular Psychiatry, 25(6), 1159–1174. https://doi.org/10.1038/s41380-019-0493-2

Tham, M. W., Woon, P. S., Sum, M. Y., Lee, T. S., & Sim, K. (2011). White matter abnormalities in major depression: Evidence from post-mortem, neuroimaging and genetic studies. Journal of Affective Disorders, 132(1–2), 26–36. https://doi.org/10.1016/j.jad.2010.09.013

Thomas, A. W., Caporale, N., Wu, C., & Wilbrecht, L. (2016). Early maternal separation impacts cognitive flexibility at the age of first independence in mice. Developmental Cognitive Neuroscience, 18, 49–56. https://doi.org/10.1016/j.dcn.2015.09.005

Thomas, A. W., Delevich, K., Chang, I., & Wilbrecht, L. (2020). Variation in early life maternal care predicts later long range frontal cortex synapse development in mice. Developmental Cognitive Neuroscience, 41, 100737. https://doi.org/10.1016/j.dcn.2019.100737

Tottenham, N., & Galván, A. (2016). Stress and the adolescent brain. Neuroscience & Biobehavioral Reviews, 70(3), 217–227. https://doi.org/10.1016/j.neubiorev.2016.07.030

Tsoory, M., Cohen, H., & Richter-Levin, G. (2007). Juvenile stress induces a predisposition to either anxiety or depressive-like symptoms following stress in adulthood. European Neuropsychopharmacology, 17(4), 245–256. https://doi.org/10.1016/j.euroneuro.2006.06.007

Tsoory, M., & Richter-Levin, G. (2006). Learning under stress in the adult rat is differentially affected by “juvenile” or “adolescent” stress. International Journal of Neuropsychopharmacology, 9(6), 713–728. https://doi.org/10.1017/S1461145705006255

Udwin, O., Boyle, S., Yule, W., Bolton, D., & O’Ryan, D. (2000). Risk factors for long-term psychological effects of a disaster experienced in adolescence: Predictors of Post Traumatic Stress Disorder. Journal of Child Psychology and Psychiatry and Allied Disciplines, 41(8), 969–979. https://doi.org/10.1017/S0021963099006460

Van Bodegom, M., Homberg, J. R., & Henckens, M. J. A. G. (2017). Modulation of the hypothalamic-pituitary-adrenal axis by early life stress exposure. Frontiers in Cellular Neuroscience, 11(April), 1–33. https://doi.org/10.3389/fncel.2017.00087

Ventriglio, A., Gentile, A., Baldessarini, R. J., & Bellomo, A. (2015). Early-Life Stress and Psychiatric Disorders: Epidemiology, Neurobiology and Innovative Pharmacological Targets. Current Pharmaceutical Design, 21(11), 1379–1387. https://doi.org/10.2174/1381612821666150105121244

Wei, L., Hao, J., Lacher, R. K., Abbott, T., Chung, L., Colangelo, C. M., & Kaffman, A. (2015). Early-Life Stress Perturbs Key Cellular Programs in the Developing Mouse Hippocampus. Developmental Neuroscience, 37(6), 476–488. https://doi.org/10.1159/000430861

Weissman, M. M., Neria, Y., Das, A., Feder, A., Blanco, C., Lantigua, R., … Olfson, M. (2005). Gender differences in posttraumatic stress disorderamong primary care patients after the World Trade Center attack of September 11, 2001. Gender Medicine, 2(2), 76–87. https://doi.org/ https://doi.org/10.1016/S1550-8579(05)80014-2

Wennström, M., Hellsten, J., Ekstrand, J., Lindgren, H., & Tingström, A. (2006). Corticosterone-induced inhibition of gliogenesis in rat hippocampus is counteracted by electroconvulsive seizures. Biological Psychiatry, 59(2), 178–186. https://doi.org/10.1016/j.biopsych.2005.08.032

Xu, W., Yao, X., Zhao, F., Zhao, H., Cheng, Z., Yang, W., … Li, B. (2020). Changes in Hippocampal Plasticity in Depression and Therapeutic Approaches Influencing These Changes. Neural Plasticity, 2020, 1–16. https://doi.org/10.1155/2020/8861903

Yang, Y., Cheng, Z., Tang, H., Jiao, H., Sun, X., Cui, Q., … Li, B. (2017). Neonatal Maternal Separation Impairs Prefrontal Cortical Myelination and Cognitive Functions in Rats Through Activation of Wnt Signaling. Cerebral Cortex, 27(5), 2871–2884. https://doi.org/10.1093/cercor/bhw121

Yehuda, R., Halligan, S. L., & Grossman, R. (2001). Childhood trauma and risk for PTSD: Relationship to intergenerational effects of trauma, parental PTSD, and cortisol excretion. Development and Psychopathology, 13, 733–753.

Zhang, H., Yan, G., Xu, H., Fang, Z., Zhang, J., Zhang, J., … Huang, Q. (2016). The recovery trajectory of adolescent social defeat stress-induced behavioral, 1H-MRS metabolites and myelin changes in Balb/c mice. Scientific Reports. https://doi.org/10.1038/srep27906

Zoladz, P. R., & Diamond, D. M. (2016). Predator-based psychosocial stress animal model of PTSD: Preclinical assessment of traumatic stress at cognitive, hormonal, pharmacological, cardiovascular and epigenetic levels of analysis. Experimental Neurology, 284, 211–219. https://doi.org/10.1016/j.expneurol.2016.06.003

Zovkic, lva B., Meadows, J. P., Kaas, G. A., & Sweatt, J. D. (2013). Interindividual variability in stress susceptibility: A role for epigenetic mechanisms in PTSD. Frontiers in Psychiatry. https://doi.org/10.3389/fpsyt.2013.00060

