## Supplemental Figures for "Juvenile exposure to acute traumatic stress leads to long-lasting alterations in grey matter myelination in adult female but not male rats"

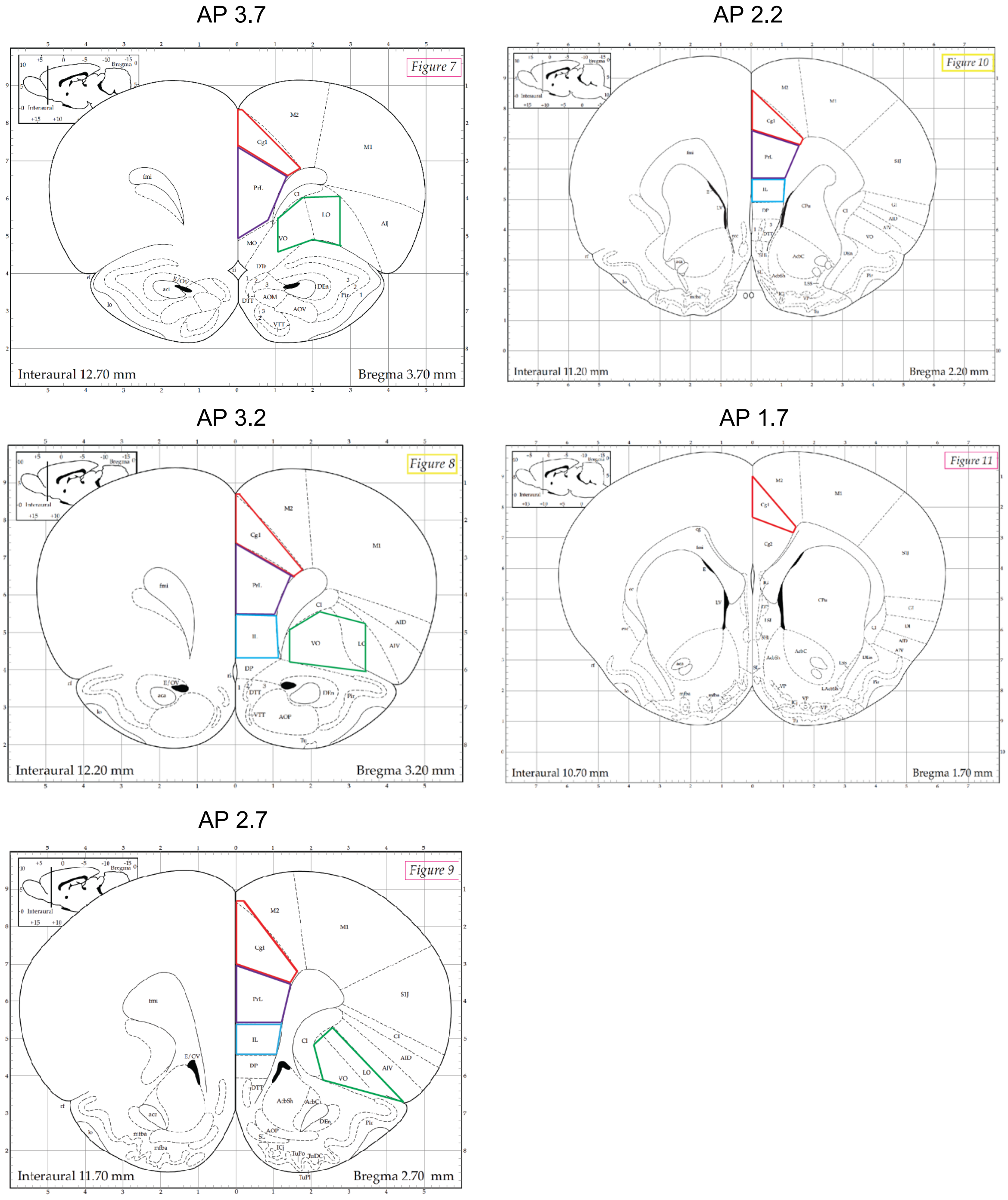


**Figure S1**. Atlas images of the five representative prefrontal cortex (PFC) regions used for immunohistochemistry. ROIs were hand drawn in ImageJ approximately as shown above. Red = Cg1, Purple = PrL, Blue = IL, Green = OFC. Coordinates are anterior-posterior (AP) from bregma (Paxinos & Watson. The Rat Brain Atlas. 1998).

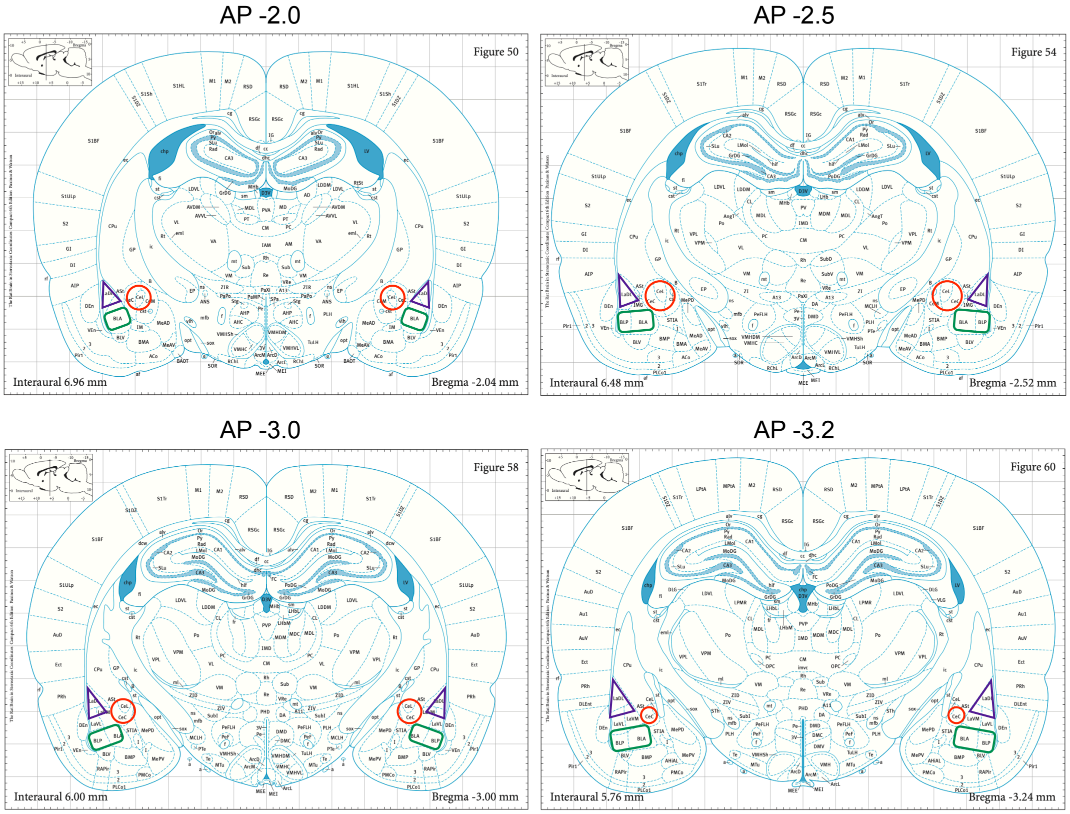


**Figure S2:** Atlas images of the four representative amygdala (AMY) regions used for immunohistochemistry. ROIs were hand drawn in ImageJ approximately as shown above. Red = CeA, Purple = LA, Green = BA. Coordinates are anterior-posterior (AP) from bregma (Paxinos & Watson. The Rat Brain Atlas. 1998).

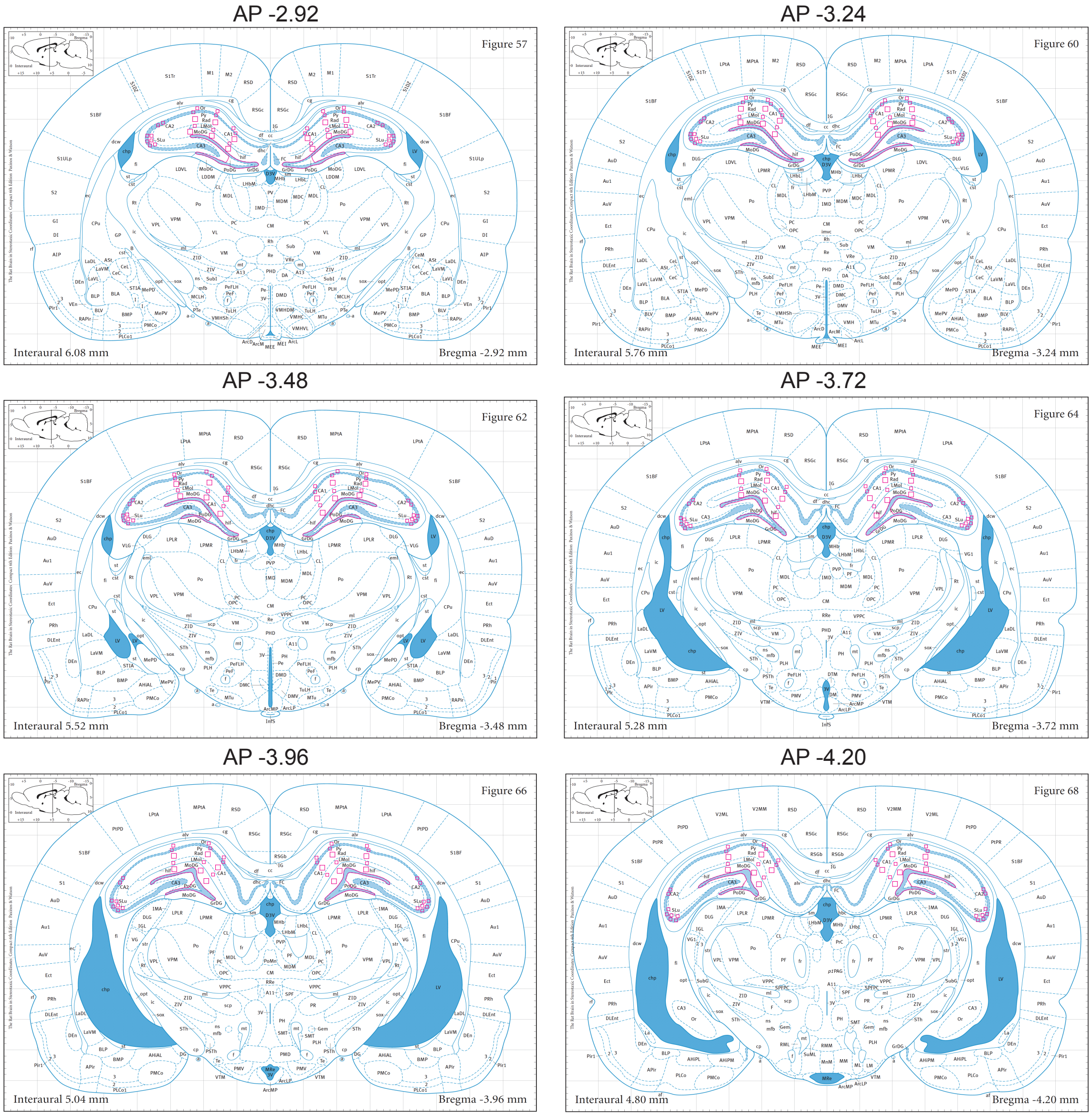


**Figure S3:** Atlas images of the six representative hippocampus (HPC) regions used for immunohistochemistry. ROIs were hand drawn in ImageJ for the Hilus and GCL layers. Squares of varying sizes were placed on all other ROIs. Coordinates are anterior-posterior (AP) from bregma (Paxinos & Watson. The Rat Brain Atlas. 1998).

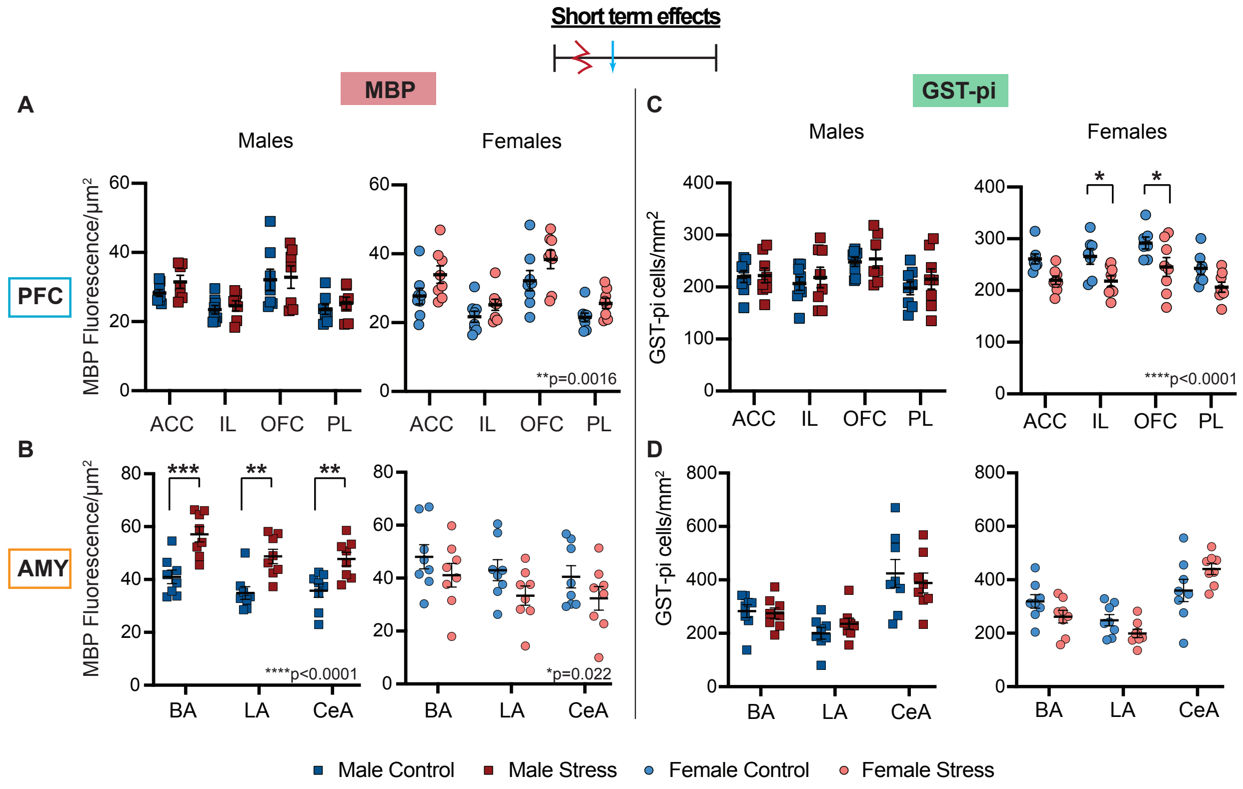


**Figure S4.** MBP and GST-pi levels within each PFC and AMY subregion, for animals tested in the short term. **A-B:** MBP changes in the **A)** PFC and **B)** AMY, and **C-D:** GST-pi changes in the **C)** PFC and **D)** AMY. The p-values of statistically significant main effects of condition are written on each graph. Statistically significant post-hoc tests are marked with asterisks (*p<0.05, ** p<0.01, ***p<0.001).

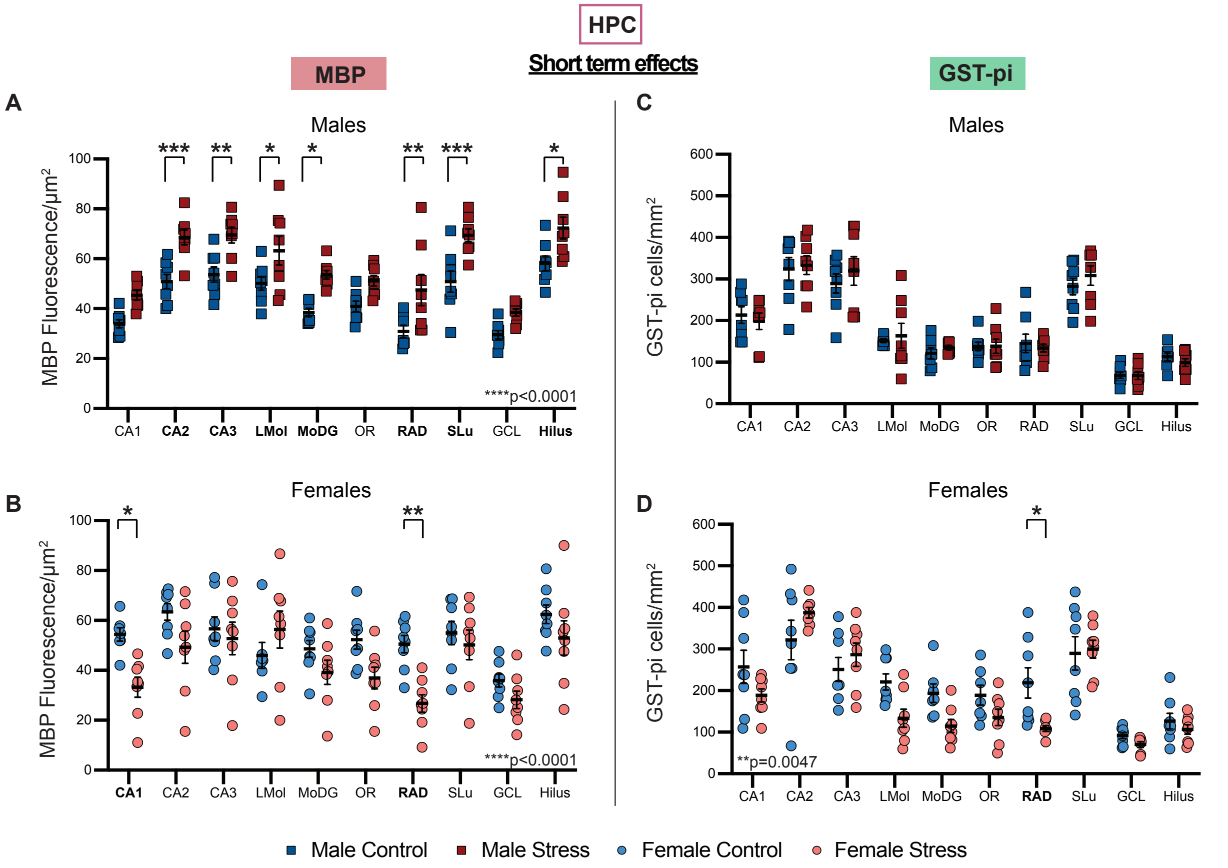


**Figure S5.** MBP and GST-pi levels within each HPC subregion, for animals tested in the short term. **A-B:** MBP changes in **A)** males and **B)** females and **C-D:** GST-pi changes in **C)** males and **D)** females. The p-values of statistically significant main effects of condition are written on each graph. Statistically significant post-hoc tests are marked with asterisks (*$p<0.05$, ** $p<0.01$, ***$p<0.001$).

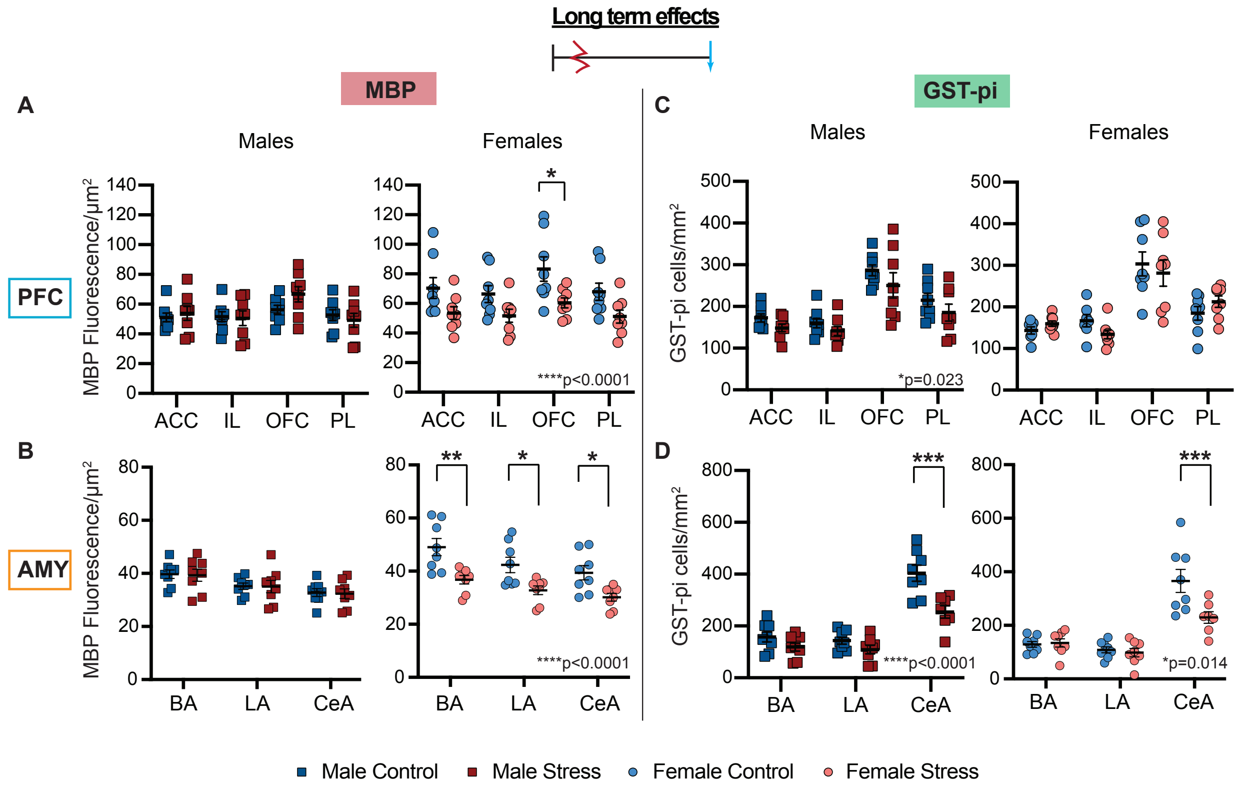


**Figure S6.** MBP and GST-pi levels within each PFC and AMY subregion, for animals tested in the long term. **A-B:** MBP changes in the **A)** PFC and **B)** AMY, and **C-D:** GST-pi changes in the **C)** PFC and **D)** AMY. The p-values of statistically significant main effects of condition are written on each graph. Statistically significant post-hoc tests are marked with asterisks ($*p<0.05$, $** p<0.01$, $***p<0.001$).

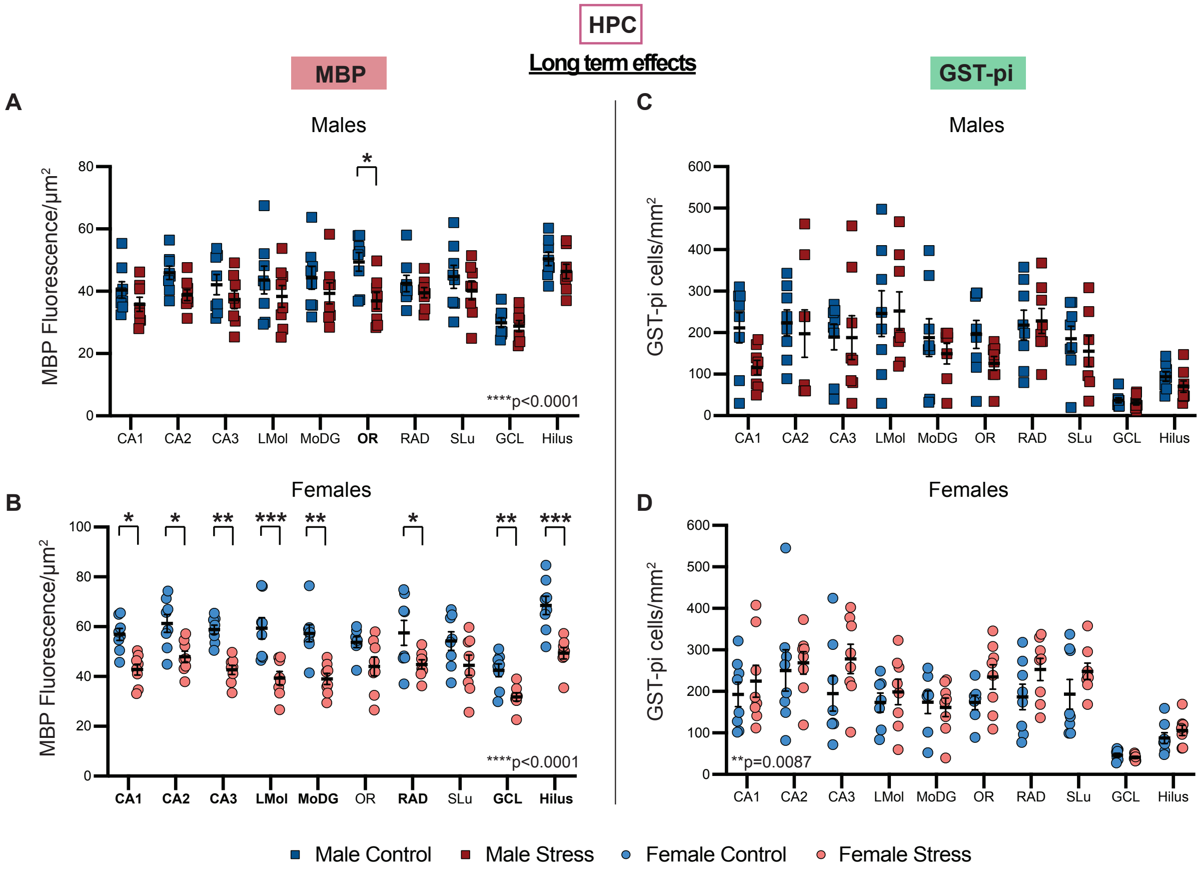


**Figure S7.** MBP and GST-pi levels within each HPC subregion, for animals tested in the long term. **A-B:** MBP changes in **A)** males and **B)** females and **C-D:** GST-pi changes in **C)** males and **D)** females. The p-values of statistically significant main effects of condition are written on each graph. Statistically significant post-hoc tests are marked with asterisks (*p<0.05, ** p<0.01, ***p<0.001).

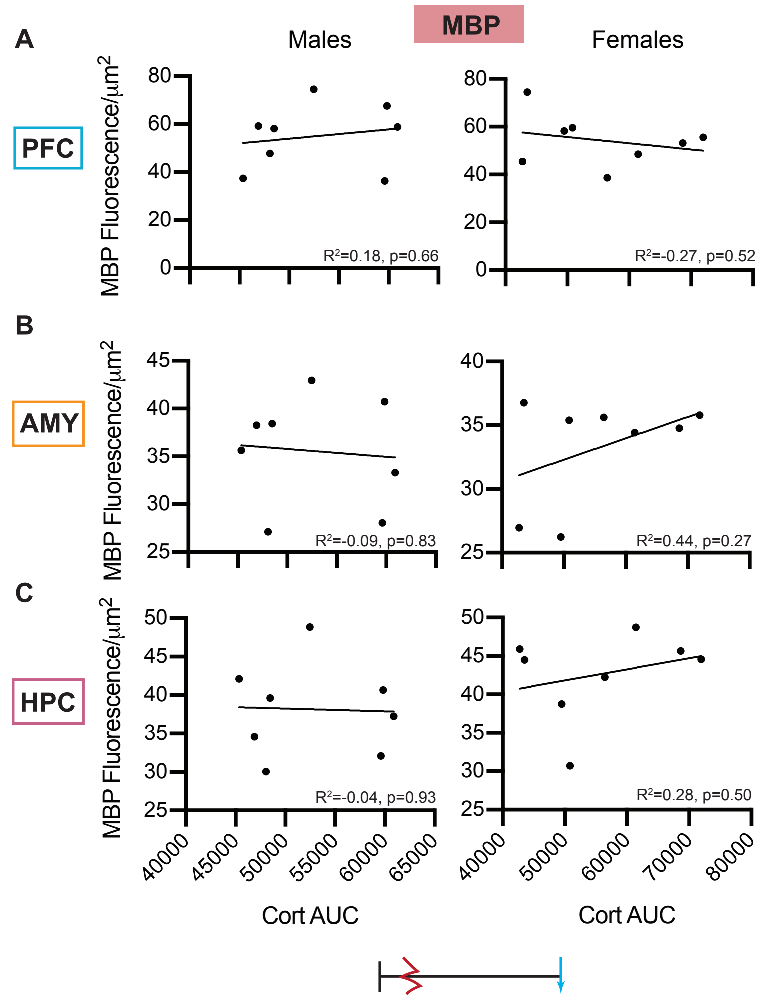


**Figure S8.** Correlations of corticosterone with long-term myelin. Pearson's correlations of MBP fluorescence intensity with area under the curve (AUC) for corticosterone in the **(A)** PFC**, (B)** AMY and **(C)** HPC of male and female rats tested at p95.

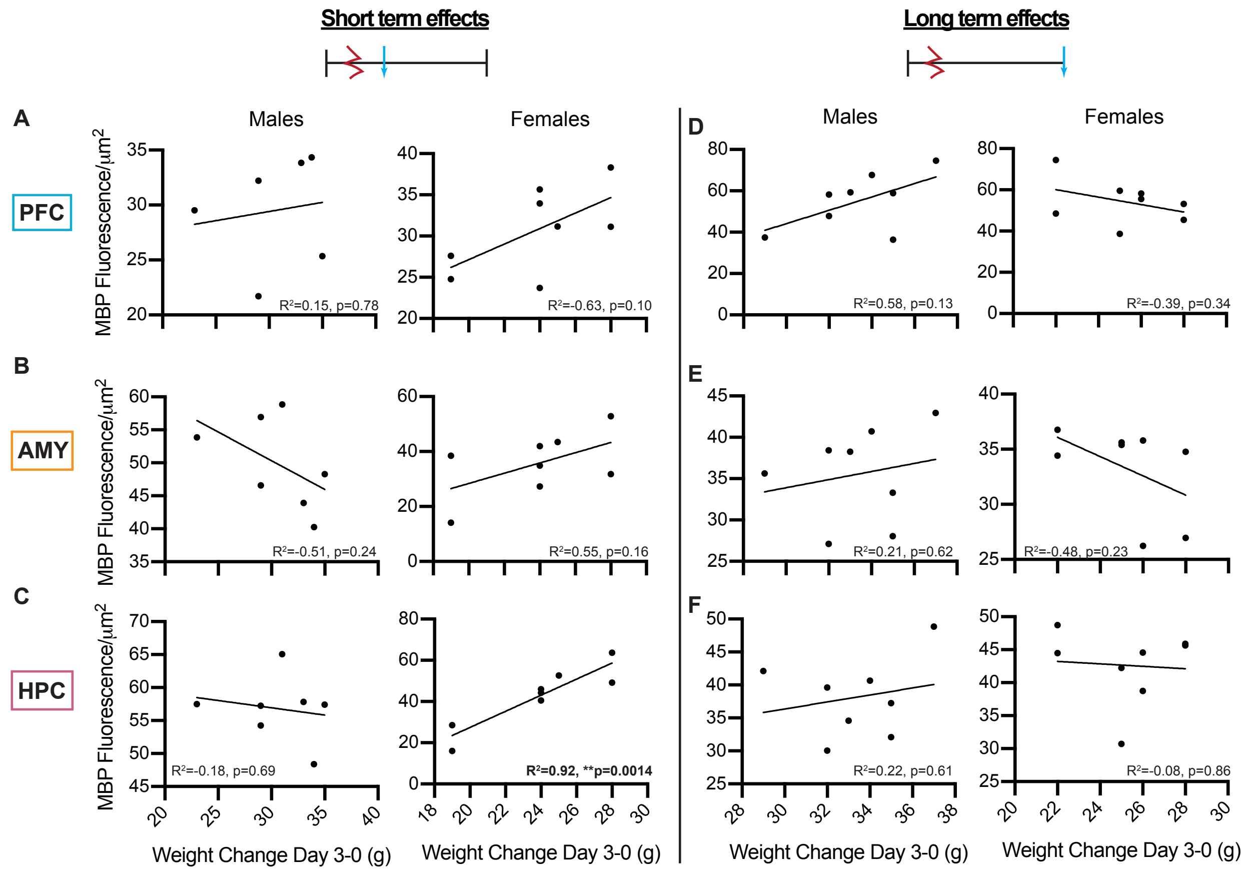


**Figure S9.** Correlations of weight change with myelin. Pearson's correlations of MBP fluorescence intensity with weight changes three days following stress exposure in the short-term **(A-C)** and in the long-term (**D-F)** in the PFC **(A,D),** AMY **(B,E)** and HPC **(C,F)** of male and female rats.
